# Ultrafast Structural Changes Decomposed from Serial Crystallographic Data

**DOI:** 10.1101/634071

**Authors:** Zhong Ren

## Abstract

Direct visualization of electronic and molecular events during chemical and biochemical reactions will offer unprecedented mechanistic insights. Ultrashort pulses produced by X-ray free electron lasers (XFELs) offer an unprecedented opportunity for direct observations of transient events as short-lived as tens of femtoseconds. This paper presents an in-depth analysis of the serial crystallographic datasets collected by Barends & Schlichting et al. (Science 350, 445, 2015) that probe the ligand photodissociation in carbonmonoxy myoglobin (MbCO), a long-serving hallmark for observing ultrafast dynamics in a biological system. This analysis reveals electron density changes that are caused directly by the formation of high-spin 3*d* atomic orbitals of the heme iron upon the CO departure and the dynamic behaviors of these newly formed orbitals in a time series within the first few picoseconds. The heme iron is found vibrating at a high frequency and developing a positional modulation that causes the iron to pop out of and recoil back into the heme plane in succession. These findings here provide long-awaited visual validations for previous works using ultrafast spectroscopy and molecular dynamics simulations. This analysis also extracts electron density variations largely in the solvent during the first period of a low frequency oscillation previously detected by coherence spectroscopy. This work demonstrates the power and importance of the analytical methods in detecting and isolating transient, often very weak signals of electronic changes arising from chemical reactions in proteins.

**Summary:** Direct imaging of ultrafast and subtle structural events during a biochemical reaction, such as a single electronic transition from one atomic or molecular orbital to another, is highly desirable but has been beyond the reach of protein crystallography. It entails the capability of observing changes in electronic distributions at both an ultrafast time scale and an ultrahigh spatial resolution (Itatani et al., Nature 432, 867, 2004). The recent developments in femtosecond serial crystallography at X-ray free electron lasers (XFELs) have brought the achievable temporal resolution within a striking distance. This paper presents the electron density map decomposed from the XFEL data that shows the remanence of several 3*d* atomic orbitals of the heme iron at an available spatial resolution although the map component is not an accurate image of the atomic orbitals. A key strategy that has enabled the findings here is a numerical deconvolution to resolve concurrent variations in a series of time-resolved electron density maps so that the electron densities influenced by an electron transfer event can be isolated as a partial change from the overwhelming presence of the bulk electrons that are not directly involved in bonding. Even at the limited spatial resolution, the subtle changes in electron distribution due to a spin crossover can be decoupled from far greater changes due to atomic displacements. Direct observations of electronic orbitals could offer unprecedented mechanistic insights into a myriad of chemical and biochemical reactions such as electron transfer in redox reactions, and formation, rupture, and isomerization of chemical bonds.

Ligand photodissociation in carbonmonoxy myoglobin (MbCO) has been a benchmark for studying ultrafast protein dynamics in a biological system. A number of studies in time-resolved crystallography have progressively improved the time resolution (from Šrajer et al., Science 274, 1726, 1996 to Barends et al., Science 350, 445, 2015). This paper presents an in-depth analysis of the serial crystallographic datasets of MbCO that Barends & Schlichting et al. (2015) contributed to the Protein Data Bank. First, a component of electron density distributions clearly shows the characteristic shape of the high-spin 3*d* orbitals reappeared at the heme iron upon the photodissociation of the CO ligand despite the limited accuracy of the orbital image due to the available spatial resolution. Second, the dynamic behaviors of these newly regained 3*d* orbitals within picoseconds after the photolysis provide long-awaited structural validation for previous spectroscopic observations and computational simulations. Specifically, the newly formed densities are oscillating with the heme iron at a high frequency of a thousand wavenumbers and developing a positional modulation during the first few picoseconds (Champion, Science 310, 980, 2005). The iron pops out of the heme plane at a few picoseconds and recoils back and pops out again afterwards. The dominant oscillation at a low frequency of several tens wavenumbers previously detected by coherence spectroscopy can be clearly resolved from the time series of electron density maps. The associated changes in electron density during the first cycle of the oscillation are largely located in the solvent rather than on the protein or heme, which suggests that the low frequency oscillations in a number of heme proteins, including MbCO, likely originate from a photolysis triggered pressure wave propagating in the solvated protein. Finally, these findings of chemical signals are isolated from coexisting thermal artifacts also by the numerical deconvolution. It is modeled in this study that the ultrashort XFEL pulses cause a transient spike of the local temperature at the heme site of hundreds of K.

## Introduction

One of the major applications of X-ray free electron lasers (XFELs) is to directly observe ultrafast protein dynamics at atomic resolution during photochemical reactions by serial femtosecond (fs) crystallography (SFX). Time-resolved experiments (Barends et al., 2015; Coquelle et al., 2017; Nango et al., 2016; Nogly et al., 2018; Pande et al., 2016; Tenboer et al., 2014) have been successfully conducted by several groups at XFELs to follow the photodissociation of carbonmonoxy myoglobin (MbCO; Fig. S1), photoisomerization of the *p*-coumaric acid chromophore in photoactive yellow protein (PYP), and the hydroxybenzylidene imidazolinone chromophore in a photoswitchable fluorescent protein (Levantino et al., 2015a). While the time resolution, that is, the ability to differentiate transient changes, is limited to tens of fs determined by the duration of the XFEL pulses, the reported time resolution for MbCO and PYP was around hundreds of fs due to some timing jitter between a short laser pump and an XFEL probe.

It is well established that a ligand bound to the heme iron helps to split its five 3*d* orbitals into two energy levels, the triply degenerate *t*2g at a lower energy and the doubly degenerate *e*g at a higher energy, so that the six 3*d* electrons of Fe(II) form three pairs occupying the lower energy orbitals, thus the net spin is zero. Upon photodissociation, the CO ligand parks at a nearby docking site, while the heme iron moves out of the heme plane towards the proximal side. The absence of a bound ligand returns the five degenerate 3*d* orbitals of the iron back to approximately identical energy levels so that they are all occupied resulting in a high-spin, deoxy iron. As the porphyrin structure becomes buckled, several side chains and helices of the globin move accordingly (Fig. 1). These light-induced structural changes are highly reproducible and have been observed by both static and time-resolved crystallography, repeatedly (Bourgeois et al., 2003, 2006; Chu et al., 2000; Kachalova et al., 1999; Schotte et al., 2003; Šrajer et al., 1996, 2001). Recently, they have been captured once again at a sub-picosecond (sub-ps) time scale in MbCO crystals using XFEL pulses (Barends et al., 2015), a major achievement in SFX. However, no significantly new structural finding is beyond those already obtained by time-resolved Laue diffraction at synchrotrons (Bourgeois et al., 2007; Clifton et al., 1997; Ren et al., 1999), which is one thousand times slower at a time resolution of 100 ps limited by the bunch length of the synchrotrons (Aranda et al., 2006; Bourgeois et al., 2006; Jung et al., 2013; Ren et al., 2012; Schotte et al., 2012).

**Figure 1.**
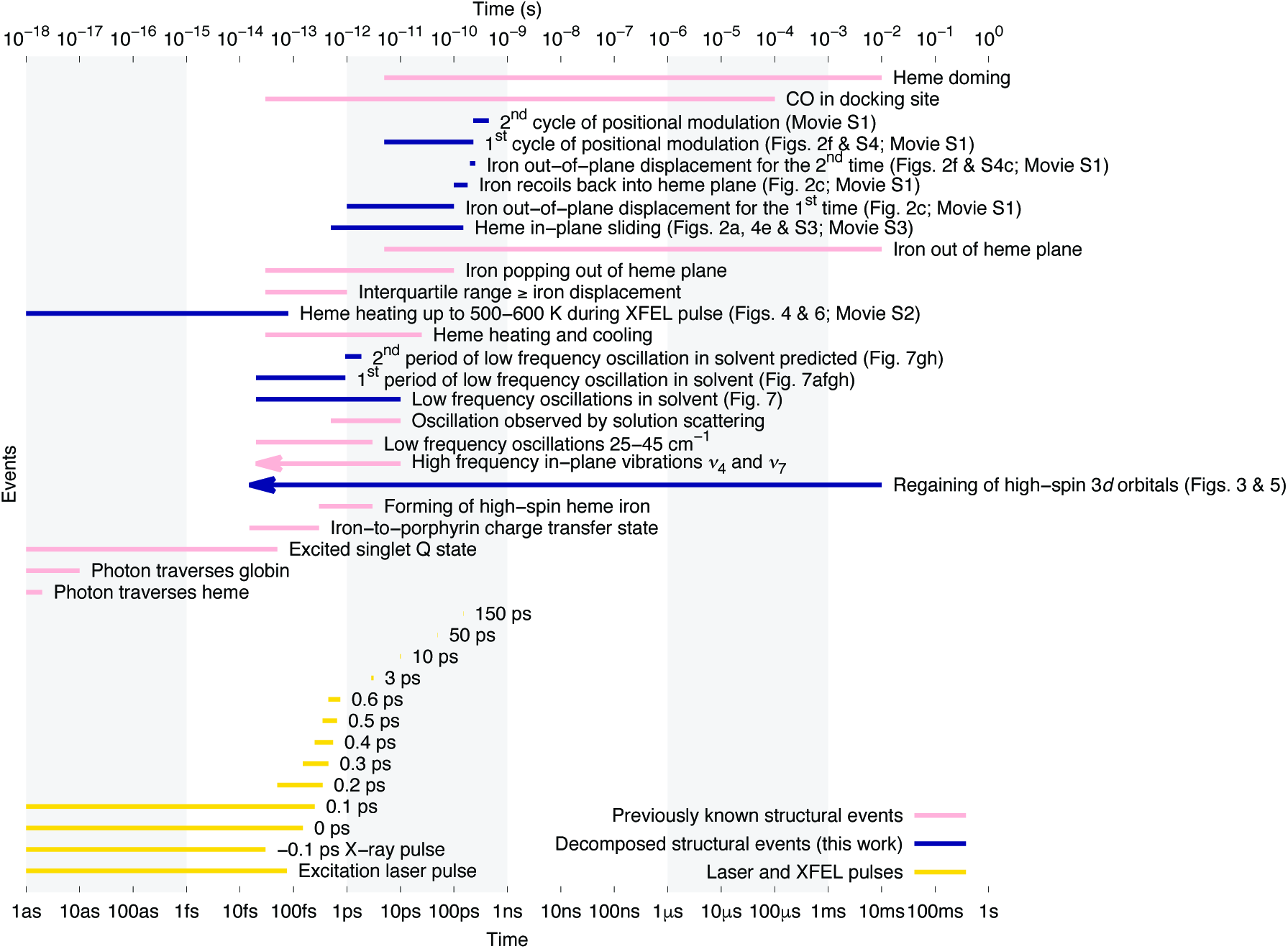
A Gantt chart summarizing structural events during photodissociation of MbCO. Previously known structural events are in pink. The excitation laser pulse and XFEL pulses are in yellow. The newly observed structural events from decomposition of the serial crystallographic datasets are in dark blue.

The signals of ultrafast structural events could be too weak to be observed clearly by the standard crystallographic analyses. In this work, I reexamine the time-resolved XFEL datasets of MbCO deposited in the Protein Data Bank (PDB; 5cmv, 5cn4-9, and 5cnb-g) by Barends & Schlichting et al. (2015) with a previously developed analytical method based on singular value decomposition (SVD; Ren, 2013a; Ren et al., 2012, 2013; Schmidt et al., 2003). This SVD analysis unravels several ultrafast events that have been identified by spectroscopy and computational studies but have not been directly observed as images of electron density distribution. The new findings along with several previously known structural events upon photolysis of MbCO are summarized in a Gantt chart (Fig. 1).

### Heterogeneous signals in dynamic crystallography

Compared to the well-established structural events of MbCO photolysis such as ligand docking and heme doming, the ultrafast signals are small in amplitude and low in population. As a result, they do not stand out at a given time point, thus went unnoticed in the conventional analyses. As demonstrated below, these weak yet consistent signals can be manifested as major components resulting from SVD when multiple datasets collected at different time points are jointly examined. This necessity for decomposition presents both a challenge and a unique opportunity for dynamic crystallography, broadly defined as experimental and analytical approaches that tightly join crystallographic datasets and their metadata (Ren et al., 2013). We have developed a numerical process of deconvolution, which is able to identify and extract the dynamic signals from a large collection of datasets acquired at varying experimental conditions such as time points and temperatures (Bandara et al., 2017; Kim et al., 2017; Ren, 2016; Ren et al., 2012; Yang et al., 2011; Zeng et al., 2015). In essence, this strategy takes advantage of the structural heterogeneity, rather than preemptively avoiding them as in the common practice of static protein crystallography. In a nutshell, dynamic crystallography consists of two major steps: first, to experimentally introduce a functionally-relevant structural heterogeneity; and second, to numerically resolve the heterogeneous conformational species as long as their compositions vary as a function of an experimental parameter (Ren et al., 2013). Here, time-resolved serial crystallography, with the variable being the ultrashort time delay, is simply a special case in the broader scope of dynamic crystallography. Therefore, the previously established methodology is largely applicable (Methods).

### Ultrafast dynamics during photodissociation of MbCO

The departure of the CO ligand from the iron porphyrin system has been said within the first 15-20 fs upon photolysis (Dunietz et al., 2003; Falahati et al., 2017, 2018). This signal is clear in the earliest time point of ∼30 fs (nominally −0.1 ps; Fig. S2). However, the photolyzed CO ligand is already observed in its docking site 3.5-4.5 Å away from the heme iron at tens to hundreds of fs (Figs. 2c and 3agj) unlike recently suggested by quantum wavepacket dynamics that the ligand departs only 0.4±0.1 Å (Falahati et al., 2018). A transient charge transfer from the iron to porphyrin subsequently taking place during 50-300 fs has been identified as a key event by resonance Raman spectra (Franzen et al., 2001). A high-spin iron is resulting in the charge transfer from the porphyrin back to the iron. The transient charge transfer state is not visible in the maps calculated from the datasets of Barends & Schlichting et al. perhaps due to a broad distribution of the transferred electron over the porphyrin. However, the larger, high-spin iron is clearly manifested in the earliest maps (Figs. 2, 3, and S2). Vibrational signals have been observed by ultrafast spectroscopy, which is attributed to a discontinuity or “seam” at the intersection of two energy landscapes of the photoexcited MbCO and the ligand dissociated product (Champion, 2005). Dissipation of such vibrational energy in the protein matrix and its solvent environment consists of at least two distinct components: localized, high frequency vibration modes of the heme at approximately one thousand wavenumbers; and global, low frequency oscillations of the protein at several tens wavenumbers (Fujisaki and Straub, 2005). This analysis provides direct visual evidences to validate both components and to delineate the low frequency oscillations in greater details (Fig. 7).

**Figure 2.**
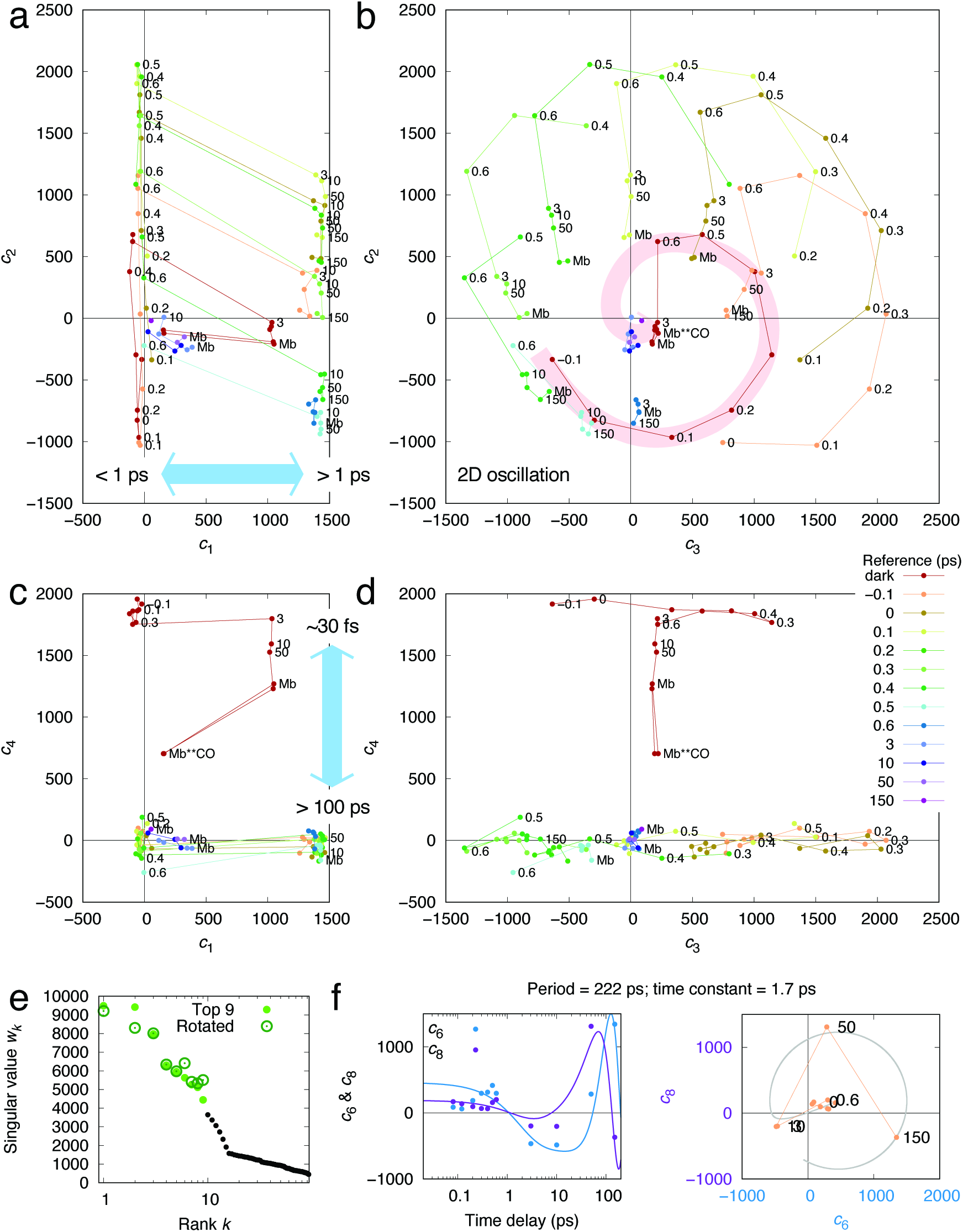
SVD of 93 difference maps around the heme site. These difference maps are divided into several series. Each series consists of the difference maps produced from subtracting the same reference (Methods and Table S1). Each of the 93 difference maps can be expressed as a linear combination of many components *c*_1_***U***_1_ + *c*_2_***U***_2_ + … after the decomposition. Among a common set of basis components ***U***_*k*_ (*k* = 1, 2, …) shared by all difference maps, also known as left singular vectors in linear algebra, the first few components are the most significant. Each of the difference maps requires a coefficient set *ck* for the linear combination. This coefficient set varies from one map to the other, that is, a function of the delay time or other metadata associate with the datasets. Each coefficient *ck* indicates how much of the corresponding component ***U***_*k*_ exists in the linear combination. Each coefficient set is derived from a right singular vector as traditionally known in linear algebra (Methods). These coefficients produced by SVD should not be confused with the Fourier coefficients used to synthesize each map. (a-d) Orthographic projections of coefficients in some of the significant dimensions. Each difference map is represented by a dot at the location that marks the coefficients associated with the major components ***U***_1_ through ***U***_4_. Difference maps in a series are connected by lines in a corresponding color. The multi-dimensional space is projected to various planes and tiled together so that any two adjacent panels from (a) to (d), horizontal or vertical, can be imagined erecting a three-dimensional subspace. (e) Singular values indicating the significance of the components are sorted in a descending order of their values before a rotation. A multi-dimensional rotation alters their values. (f) Coefficients *c*_6_ and *c*_8_ of the -*F*_−0.1_ series in yellow are least-squares fitted with a two-dimensional oscillation (Methods).

**Figure 3.**
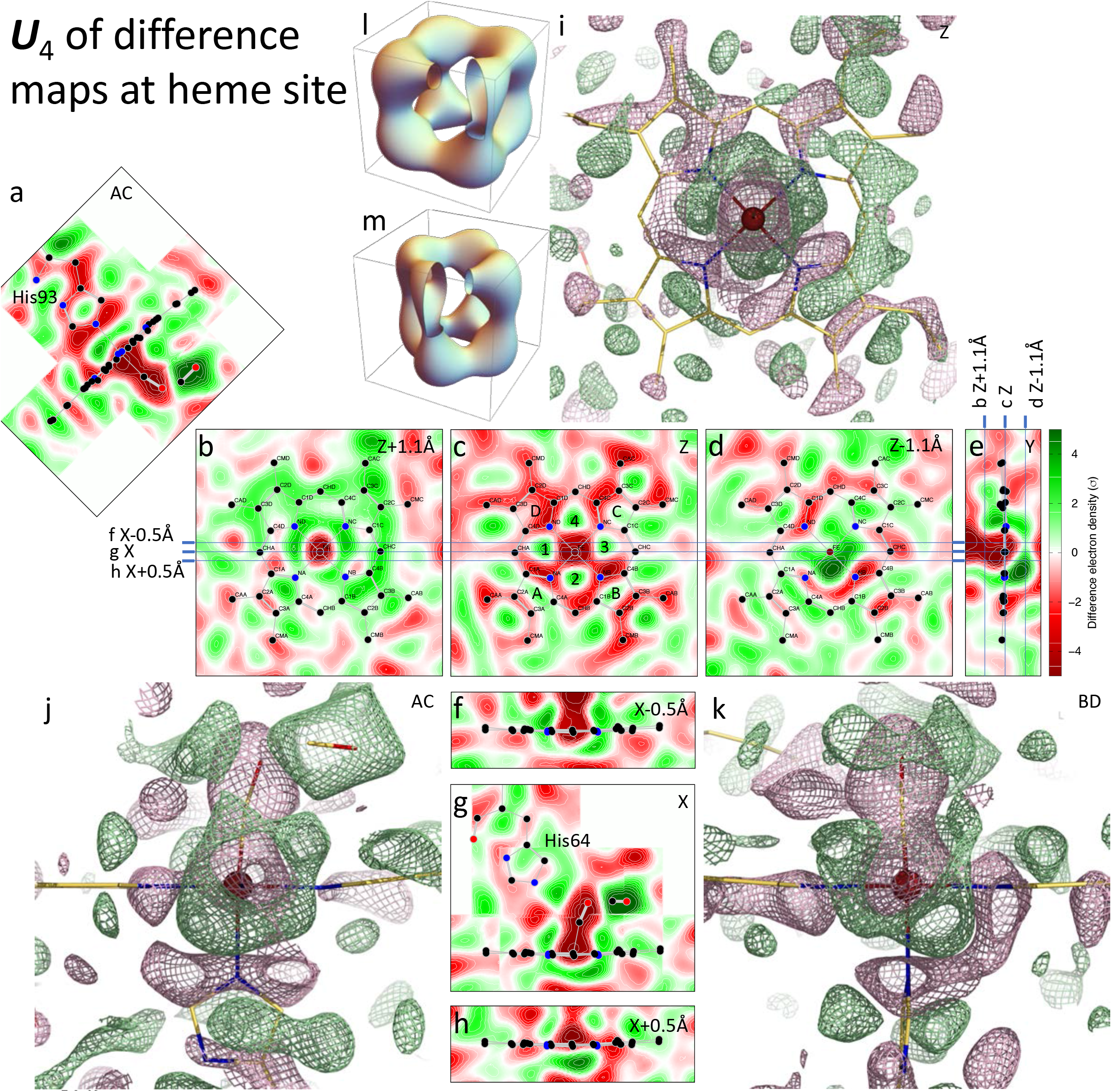
Ultrafast formation of high-spin 3*d* orbitals of the heme iron. Each component resulting from SVD, that is, a left singular vector, can also be presented as a difference map. The fourth component ***U***_4_ of the difference maps captures a cubic shaped positive density network. It is rendered as a difference map in two styles, two-dimensional cross sections and three-dimensional contours. The corresponding coefficient *c*_4_ is plotted in Fig. 2cd. (a-h) Images of two-dimensional cross sections through the component difference map ***U***_4_. Cross sections defined in Fig. S1 are labeled in the top right corner of each panel. Non-centric cross sections are marked to indicate small offsets from the centric cross sections. The structure of the heme, CO ligand in both bound and docked positions, and some side chains are also projected onto the images. C, N, O, and Fe atoms are in black, blue, red, and rusty dots, respectively. The imidazole rings of the distal His64 and proximal His93 are approximately coplanar with *X* and AC cross sections, respectively. Two propionic acid side chains of AD rings extend to the left in (bcd). But they are not coplanar with these cross sections. (i-k) Three-dimensional mesh contours superimposed over a structure are traditionally rendered to display electron density maps. Green and red meshes are positive and negative contours at ±7*σ*, respectively. (l and m) Theoretical contours of probability distributions of electrons in 3*d* orbitals of iron. The combined probability distributions of 3*d_xz_* and 3*d_yz_* with 3*d_xy_* (l) or with 3*d*_*x*^2^-*y*^2^_ (m) are both in the shape of a cube with the largest probability at the eight corners. A 1/16 cutaway in each contour shows the cross sections. This characteristic shape of electron distribution is observed in ***U***_4_ of the difference maps.

The most observed high frequency modes range from hundreds to more than a thousand wavenumbers (Armstrong et al., 2003; Kitagawa et al., 2002; Mizutani, 1997). The corresponding vibrational periods of these high frequency modes of 24-50 fs are comparable to the XFEL pulse duration. Hence an individual cycle of such vibrations is beyond what can be traced out by the current XFEL datasets. Nevertheless, the averaged effect of these high frequency vibrations is detectable (Figs. 2 and 3) in the XFEL datasets of Barends & Schlichting et al. Amplitude modulation and damping of the high frequency mode *ν*7 was simulated with a period of 350 fs and a time constant of 1.3 ps (Barends et al., 2015). There are not sufficient time points in the existing datasets to determine these amplitude modulation and damping. However, a slower modulation of the high frequency modes is clearly present in these datasets (Figs. 2 and S4; Movie S1). These in-plane modes are thought to be coupled to out-of-plane motions of the heme and its iron (Nagy et al., 2005), which take place within a fraction of ps and continue to develop non-exponentially for another hundreds of ps according to molecular dynamics simulations (Henry et al., 1985; Kuczera et al., 1993). Recent simulations (Barends et al., 2015; Falahati et al., 2017) suggested that many transient species, including those with iron in or out of the plane, are mixed together before 1 ps. It was also suspected that the iron may recoil back into the heme plane after its initial drop out of the plane (Gruia et al., 2008). This analysis of the XFEL datasets finds that the out-of-plane displacement of the iron takes place simultaneously with an in-plane shift of the porphyrin (Movie S3). There is clear visual evidence to show that the first out-of-plane move peaks at a few ps and that the recoil and the second drop take place afterwards (Figs. 2cf and S4c; Movie S1). The signal associated with the iron displacement lingers on for many μs before the ligand rebinding as captured by the time-resolved Laue diffraction (Šrajer et al., 2001).

**Figure 4.**
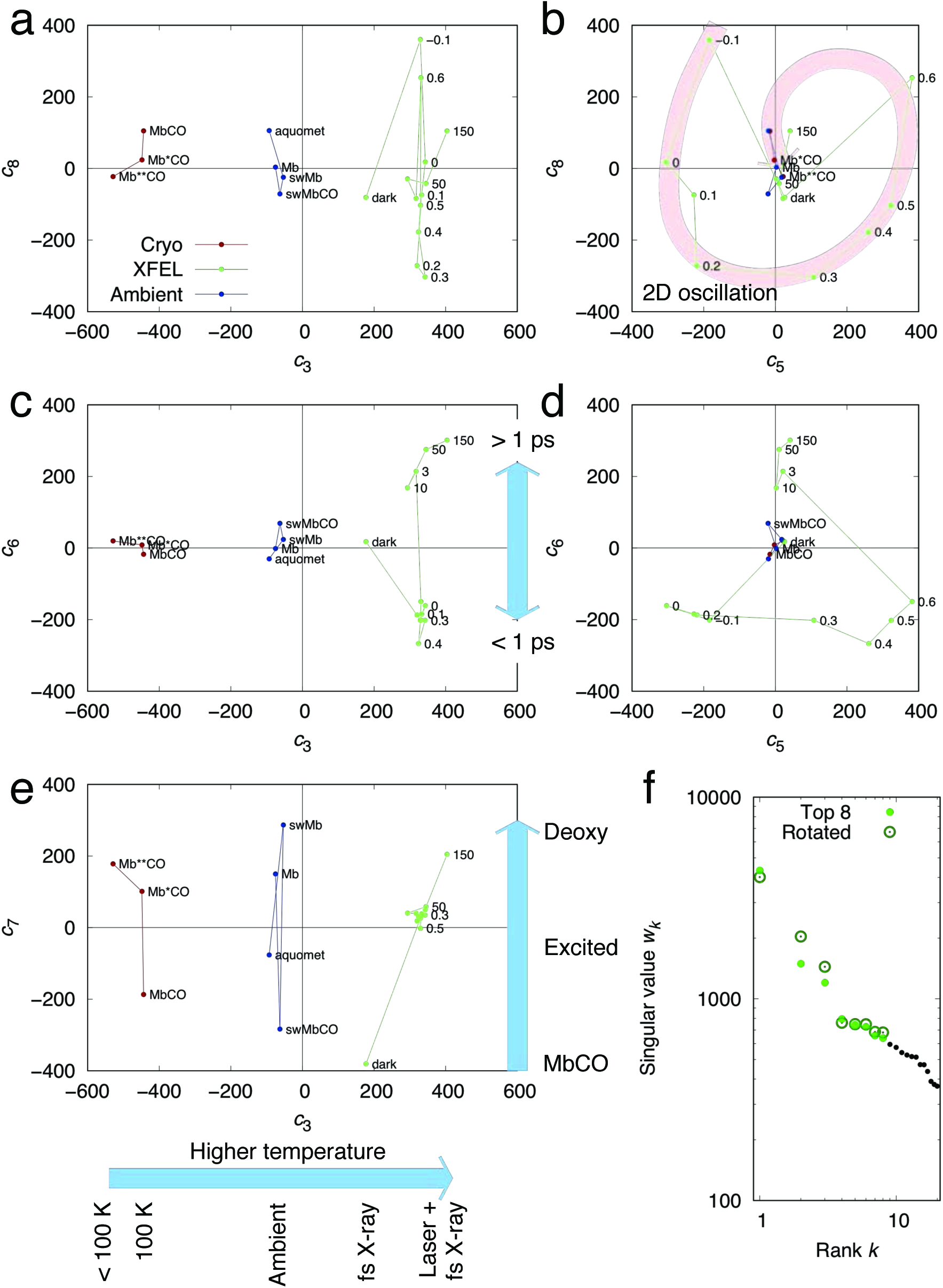
SVD of 20 simulated annealing omit maps (SAOMs) around the heme site. The first component is the average of all omit maps (Fig. S5). (a-e) Each omit map is represented by a dot at the location that marks the coefficients associated with the major components. SAOMs derived from the structures at cryo and ambient temperatures are in red and blue, respectively. SAOMs from the XFEL datasets are in green. The multi-dimensional space is projected to various planes and tiled together so that any two adjacent panels from (a) to (e), horizontal or vertical, can be imagined erecting a three-dimensional subspace. (b) Coefficients *c*_5_ and *c*_8_ show a damped two-dimensional oscillation as marked by the curved arrow. (f) Singular values indicating the significance of the components are sorted in a descending order of their values before a multi-dimensional rotation.

At a low frequency region, the dominant oscillations range between 25 and 45 cm^-1^, corresponding to a period around 1 ps (Gruia et al., 2008; Rosca et al., 2002). Such a low frequency oscillation is within the reach of the sub-ps time resolution achieved with XFEL pulses. Therefore, each individual cycle of the low frequency oscillation should be, and indeed is, visible in the datasets of Barends & Schlichting et al. Here I demonstrate that such an oscillation occurs more in the solvent channels than in the protein or the heme at a low frequency comparable to those detected by spectroscopy (Fig. 7). This analysis supplies the long missing visual evidence to support such a low-frequency oscillation in a solvated protein.

### Imaging the geometric shapes of atomic orbitals

The shape of a *d* orbital of copper was first experimentally imaged by Zuo et al. in their difference electron density map (Humphreys, 1999; Zuo et al., 1999), which is not, however, without controversy (Mulder, 2011; Scerri, 2000). Nevertheless, experimenters continued to draw parallels between captured images and atomic or molecular orbitals, sometimes with the analytical power of digital reconstructions. Molecular orbital tomography constructs images of highest occupied molecular orbital from many projections of aligned molecules (Itatani et al., 2004; Ogilvie, 2011; Schwarz, 2006; Stapelfeldt, 2004; Vozzi et al., 2011). Both the highest occupied and lowest unoccupied molecular orbitals of flat molecules, such as pentacene, were scanned using scanning tunneling microscopy (Chaika, 2014; Patera et al., 2019; Repp et al., 2005). Field-emission electron microscopy and angle-resolved photoemission spectroscopy can directly capture or indirectly reconstruct orbital images or transformation of orbital configurations (Mikhailovskij et al., 2009; Puschnig et al., 2009). More recently, a photoionization experiment showed the nodal structures of the atomic orbital of excited hydrogen atoms (Stodolna et al., 2013). A charge-density analysis at an ultrahigh resolution of 0.48 Å showed iron 3*d* and sulfur 3*p* densities (Hirano et al., 2016). In all these achievements, the experimentally captured electron densities with their unique shapes are directly influenced by electrons occupying specific orbitals. Therefore, in this paper, the images and motion pictures of difference electron densities with their characteristic nonspherical shape are attributed to a superposition of several atomic orbitals and their dynamics. However, this is not to say that the accurate image of an atomic orbital has been captured, which is not a proper task for protein crystallography, given the limit in spatial and temporal resolutions.

## Results

### Decomposition of difference maps and simulated annealing omit maps

To analyze the time-resolved serial crystallographic data of horse heart (hh) MbCO, one dark and 12 light datasets are obtained from PDB (Fig. 1; Barends et al., 2015), among which three light datasets collected at the shortest time delays of 0 and ±0.1 ps span both sides of time 0 that is centered on the excitation laser flash of 150 fs. These datasets are expected to capture ultrafast signals at delay times as short as a few fs to tens of fs. However, these ultrafast signals are also inevitably mixed with signals at longer delays as far as the X-ray pulses stretch (Haldrup et al., 2011). These sub-ps time points were taken at the same controlled time delay of 0.5 ps. But the diffraction images were binned according to the timestamp records from an experimental timing tool (Bionta et al., 2011, 2014), which produced a number of short delays before 1 ps. Several longer time points ranging from 3 to 150 ps are also included in this analysis (Fig. 1) along with a deoxy hhMb dataset collected from a synchrotron beamline at 7°C, which represents the end product of the MbCO photolysis, and two hhMb structures (1dws/t) in the photolyzed state captured by cryo-trapping (Chu et al., 2000).

Two types of electron density maps, namely difference Fourier map and simulated annealing omit map (SAOM), are produced and subjected to the SVD analyses separately. Each type carries its own advantage (Methods). While difference Fourier maps are sensitive for detecting weak signals, they are also highly susceptible to systematic errors that often overwhelm the signal. It is critically important to choose a proper reference dataset, which is preferably obtained under the identical experimental condition, if not from the very same crystal as the light dataset. An alternative is to use SAOMs, where a small portion of a protein structure is set aside from structure refinement using the simulated annealing algorithm. In such omit maps, the electron densities within the omitted region shall not be biased by the protein model, thus are considered as the most authentic representation of the experimentally observed electron densities (Hodel et al., 1992). A collection of SAOMs within a common omitted region, although less sensitive to subtle changes compared to difference maps, permits a joint analysis of maps derived under very different experimental conditions, which is usually difficult with difference maps.

Difference Fourier maps are synthesized using a Fourier coefficient set of *F*_light_-*F*_reference_ combined with the best possible phase set. While *F*_reference_ is typically the structure factor amplitudes from a dark dataset (5cmv), it is important that a light dataset at a given time point is also used as a reference (Table S1) to reveal changes with respect to that specific time point (Methods). Therefore, a collection of difference maps produced with the same reference dataset constitutes a time series that show continuous changes from the reference time point and on. For example, -*F*_0_ series refers to the difference maps with the reference dataset at the nominal 0 ps. For omit maps, the heme and several side chains within its immediate vicinity are omitted to produce SAOMs of the ligand binding site (Methods). A few static structures determined at room temperature of sperm whale (sw) Mb in the aquomet (1bz6), deoxy (1bzp), and CO-bound (1bzr) forms are also included (Kachalova et al., 1999), along with three hhMb structures (1dwr/s/t) (Chu et al., 2000) as control data points in this joint analysis. The resolutions of these static datasets are truncated to 1.75 Å to match the XFEL datasets.

Altogether, a total of 93 difference maps (Table S1 and Fig. S2) and 20 omit maps are obtained for two independent SVD analyses. The electron densities are masked around the heme site unless stated otherwise (Methods). In this work, the significant signals are identified by the top nine components from SVD (Fig. 2e). A numerical procedure called rotation in the SVD space plays an important role to extract chemically sensible components from these (difference) electron density maps (Henry and Hofrichter, 1992; Ren, 2013a, 2013b, 2016). Proper SVD rotations reveal the correlation between the core data, i.e., electron densities, and their metadata, i.e., delay time, sample temperature, and other experimental parameters (Methods). Given their highly dynamic nature, the ultrafast signals due to multiple causes are often entangled before suitable rotations are found in the multi-dimensional SVD space (Methods). Using this decomposition and rotation strategy, I identify signals that reveal the electronic redistributions between high and low spin of the iron orbitals at the available spatial resolution.

### Molecular image of low-spin to high-spin crossover

The ultrafast signals within several fs to a few tens of fs are manifested in the fourth component, denoted as ***U***_4_, of 93 difference maps. The component ***U***_4_ exists only in the map series with the dark dataset as reference (red, -*F*_dark_ series in Fig. 2). That is to say, all signals in this component have occurred by the time of the very first time point at ∼30 fs (nominally −0.1 ps), therefore the map series -*F*_−0.1_ shows the cancellation of this component (orange series in Fig. 2). Some of these signals start to decay slightly before 1 ps and diminish significantly after 3 ps (Fig. 2cd). The corresponding map of ***U***_4_ reveals the previously well-established events arising from the CO docking, out-of-plane motion of the iron, and an uneven heme doming upon photolysis of MbCO (Fig. 3). The negative densities associated with the pyrrole rings and their propionic acid side chains increase in intensity from ring A to B to C to D (Fig. 3c). Positive densities are found at a cross section 1.1 Å above the heme plane towards the distal side (Fig. 3b).

More remarkably, ***U***_4_ captures some hitherto unseen signals characterized by a cubic-shaped network of positive densities that cage the iron atom. To facilitate a more concise description, a cross section of *Z* = 0 is defined as the heme plane and the cross section of *X* = 0 is the dividing plane between two propionic acid side chains, from which the third orthogonal cross section of *Y* = 0 can be derived (Fig. S1). The eight corners of the positive cube are oriented along the *X* and *Y* cross sections. Strong positive densities appear in the middle of four six-membered rings encircled by adjacent pyrroles, the connecting methane bridge, and the heme iron (marked 1-4 in Fig. 3c). These positive features reach their four minima when they cross the heme plane. These positive densities become more prominent above and below the heme plane (Fig. 3). The eight corners of this cube are located at 1.7±0.2 Å away from the iron. This cubic-shaped network is asymmetric with a notable skew towards CD rings (Fig. 3fh). Meanwhile, the iron skews in the opposite direction towards the AB rings with an out-of-plane displacement towards the proximal side (Fig. 3e). But the major features of this cube seem more symmetric in the *Y* cross section.

Similar features are also evident in the seventh component ***U***_7_ from an independent SVD analysis of 20 SAOMs. It is clear that the seventh dimension after a proper rotation (Methods) represents the reaction coordinate of the ligand dissociation. Mb structures along this reaction coordinate form a lineup on *c*_7_ dimension regardless of sample temperatures: from the ligated structures of MbCO, to the time-resolved structures, to several photoexcited species including the static structures Mb*CO and **CO, eventually to deoxy Mb (Fig. 4e). ***U***_7_ of the omit maps contains the usual signals of ligand dissociation, such as strong negative densities on the ligand indicating dissociation and a positive layer on the distal side of the heme indicating its doming (Fig. 5), except that the quality of these signals are not as good as those from difference maps because the SAOMs are derived from datasets in very different experiments. Despite the systematic errors, the seventh component ***U***_7_ of the omit maps again clearly shows the cubic network of positive densities around the iron. This cubic network also skews towards the CD rings of the heme. This is a reoccurrence of the same signals shown in ***U***_4_ of the difference maps (compare Figs. 3 and 5). The eight corners of the positive cube are 1.9±0.2 Å away from the iron.

**Figure 5.**
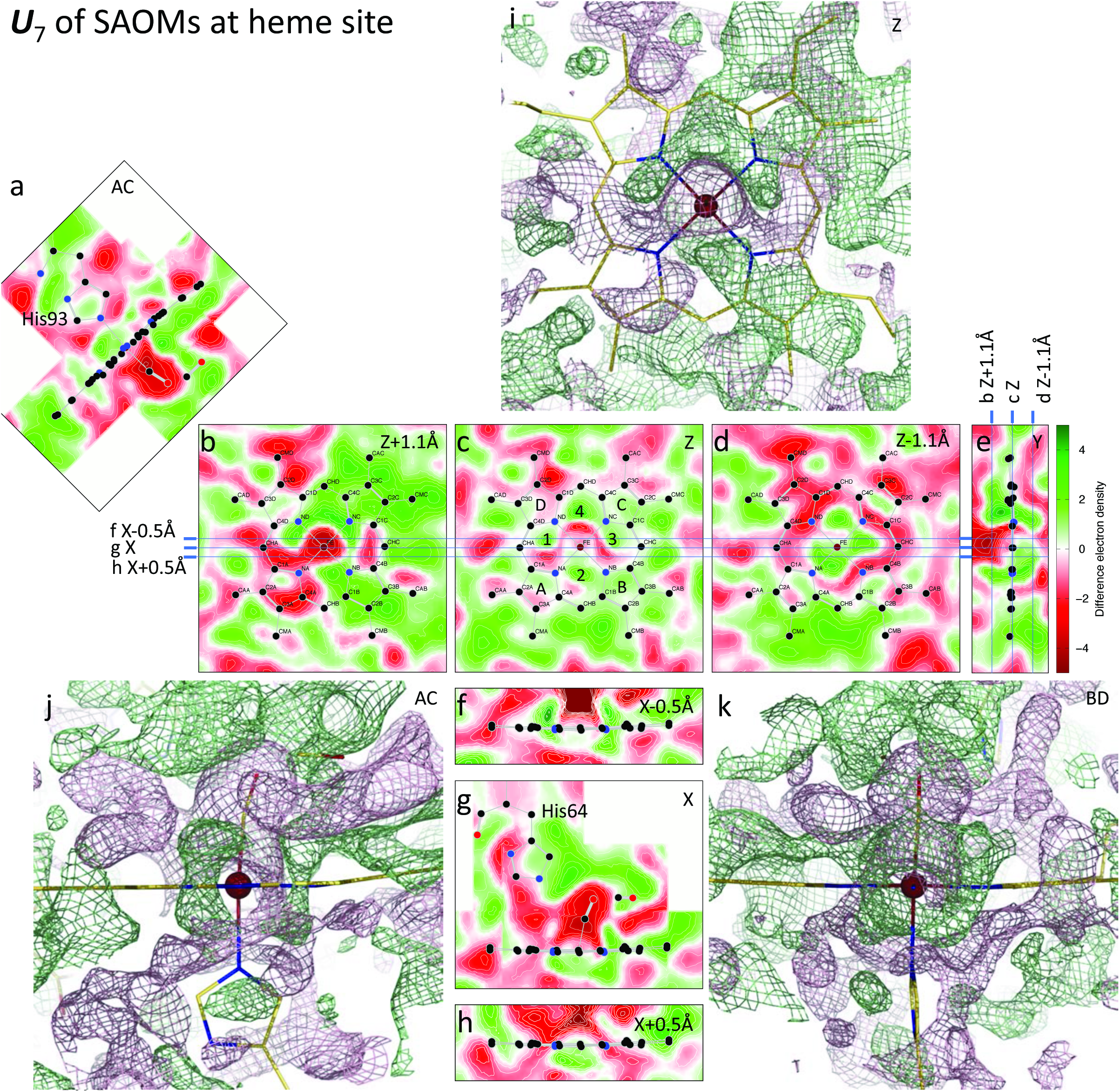
The seventh component of SAOMs correlated with photolysis at all temperatures. The coefficient corresponding to ***U***_7_ of SAOMs increases as the reaction of ligand dissociation proceeds from MbCO to deoxy Mb in static, cryo-trapping, and time-resolved XFEL experiments (Fig. 4e). The hypothetical transition from MbCO to deoxy Mb at 0 K is shown in Movie S3. (a-h) Cross sections through the seventh component ***U***_7_. A cubic network of positive densities is marked 1-4 in (c). (i-k) Three-dimensional meshes of this component in green and red are contoured at ±15*σ*, respectively. See Fig. 3 legend for more detail.

It is clear that the positive densities arranged in the cubic network around the iron, either in the difference maps or in the omit maps, cannot be easily attributed to accidental noise or systematic errors such as thermal effect or Fourier truncation error (see Discussion). They originate from the intrinsic signals induced by light and coexist with the previously seen signals of ligand dissociation. While multiple interpretations of these distinct cube-shaped signals may apply, this positive network clearly indicates a larger, high-spin iron as an immediate overall effect upon photolysis. Here I further attribute the distinct shape of the positive feature to the instantaneous regaining of the high-spin 3*d* orbitals of the heme iron. However, the observed cubic densities here are not an accurate image of a probability distribution of a 3*d* electron. They are resulted from a net gain of electron densities as a fully ligated iron transitions to the high-spin deoxy state when several 3*d* orbitals are reoccupied by unpaired electrons. Calculations of the combined probability distributions of the textbook 3*d_xz_* and 3*d_yz_* with either 3*d_xy_* or 3*d_x_^2^_-y_^2^* orbitals show features highly similar to the observed cubic-shaped map (Fig. 3lm). The strongest electron densities are located at eight corners of the cube where the lobes of 3*d* orbitals intercept. It is entirely possible that the observed cubic densities result from not only a superposition of several 3*d* orbitals but also an average of multiple electronic configurations. See Discussion below. Such orbital changes have not been captured previously in protein crystals by X-ray crystallography.

However, the loss of the paired low-spin electrons in the ligated state is not clear in the negative densities. Presumably, the loss of the CO ligand and the drop of the iron out of the heme plane cause the displacement of many more electrons than the low-spin electrons. Negative densities in the component ***U***_4_ are overwhelming. At the available spatial resolution, the loss of the low-spin electrons is not separated from the main events upon photolysis.

Interestingly, the signals in ***U***_4_ of difference maps decay after a few ps leading up to the static structures Mb*CO, **CO, and the deoxy Mb (Fig. 2cd). Given in-plane vibration modes (Abe et al., 1978; Nagy et al., 2005) with time periods comparable to the XFEL pulse duration, the averaged electron density between vibrating atoms is expected to increase, as shown in the six-membered rings. The observed decay of ***U***_4_ can be interpreted as the damping of these vibrations (see below). These in-plane vibrations trigger the iron to pop out of the heme plane (Franzen et al., 1995). However, it remains unclear from this observation which, or a combination of which, vibration modes plays the dominant role. Importantly, this decay does not reach zero even for the static deoxy Mb (Fig. 2cd), which suggests that the above interpretation of the cubic network of positive densities as the consequence of the spin crossover largely holds, since the cubic network cannot be fully explained by high frequency vibrations. This decay of ***U***_4_ after a few ps is also an evidence to support the model of iron recoiling back into the heme plane (Gruia et al., 2008), which has to be discussed with the top component ***U***_1_ below.

### Development, modulation, and damping after 1 ps

If the cubic network of positive densities is indeed influenced by the newly formed 3*d* orbitals, the dynamic behaviors of these high-spin electrons should be visible in the time points shortly after the initial formation. The observed evolution of the cubic network provides the necessary validation. Along the dimension *c*_1_ in the SVD space of 93 difference maps, all time points after 1 ps, including the deoxy Mb (5d5r, representing a time point of ∞), are well separated from the sub-ps time points around *c*_1_ = 0 (Fig. 2ac). Therefore, the first component ***U***_1_ of the difference maps indicates a major development after 1 ps and represents more permanent changes afterwards. The cubic positive densities in ***U***_4_ have no sign in ***U***_1_. Among the major features captured by ***U***_1_ (Fig. S3), the most noticeable is a rather skewed displacement of the iron towards the AB rings of the heme. In the meanwhile, the motion of His93 corroborates the skewed displacement of the iron (Fig. S3ae). It is also shown below that the entire heme moves in its plane towards the AB rings as well. Such long-lived signals in ***U***_1_ from 1 ps and on are in good agreement with a wide distribution of individual trajectories and interquartile range of iron displacement predicted by molecular dynamics simulations (Barends et al., 2015; Henry et al., 1985; Kuczera et al., 1993). However, the cryo-trapped structures, Mb*CO and **CO, are two exceptions. They exhibit very little characteristics in ***U***_1_ (Fig. 2ac), even though they presumably represent long time delays. It can be reasoned that the skewed displacement of iron requires concerted motions in the proximal His93 and helix F, which are evidently forbidden at cryogenic temperatures.

Both ***U***_1_ and ***U***_4_ feature the signal of the iron out-of-plane displacement (Figs. 3e and S3c). Therefore, the large positive coefficients *c*_1_ and *c*_4_ occurring simultaneously around several ps indicate that the iron pops out of the heme plane most at this time but only momentarily. The later decay of the fourth component ***U***_4_ records a recoiling of the iron back into the heme plane around 100 ps (Movie S1). Another major signal revealed after 1 ps is a modulation of the new formation of 3*d* orbitals indicated by the cubic-shaped positive densities around the iron. This is a positional modulation manifested in the sixth and eighth components of the difference maps. While the corresponding coefficients *c*_6_ and *c*_8_ are nearly constant in the sub-ps range, they start to ramp up and vary at a few ps. This variation can be most easily modeled as a two-dimensional oscillation at a period of 220±20 ps given the limited time points (Fig. 2f). Additional observations in the future may update this positional oscillation. The ***U***_6_ component features a distorted network of positive densities around the heme iron (Fig. S4) similar to ***U***_4_ but with an inclination towards ring B. In ***U***_6_, the strong negative density on iron is skewed towards the proximal side in contrast to ***U***_4_ (compare Figs. 3e and S4c). The positive peak on the proximal side of the iron is also extremely skewed (Fig. S4c). In a linear combination sense, ***U***_6_ and ***U***_8_ can be interpreted as variations that are combined into ***U***_4_ resulting in an oscillation of a period at a few hundreds of ps (Fig. 2f). Specifically, the cubic-shaped densities lean towards ring D from 0.5 to 3 ps, return to the mean position after 50 ps, then sway towards ring B in the opposite direction (Fig. 2f and Movie S1). Near the end of the first oscillation, the iron again pops out of the heme plane for the second time and severely skews towards AB rings (Movie S1). These observations of the iron motions support the previous model of iron recoiling (Gruia et al., 2008). Such oscillating behavior suggests modulation and damping of the vibrational modes. In other words, the cubic network of positive densities is not only observed in the earliest time points. The motions of this network are captured in later time points as well. This 220-ps oscillation is largely along BD cross section (Fig. S1), while damping of the vibrational amplitude was only recorded up to 150 ps thus insufficient to depict a complete picture. Two other more significant components ***U***_2_ and ***U***_3_ also show an oscillation of a distinct type as discussed below (Fig. 2b).

### Vibration induced by XFEL pulses and instantaneous local temperature jump

An independent SVD analysis of 20 simulated annealing omit maps (SAOMs) masked around the heme site (Methods) offers a direct comparison of the XFEL data with previous synchrotron data. Among the top eight components of the SAOMs, the first component reveals a well-defined molecular image of the heme and its surrounding (Fig. S5); and the second and fourth components apparently account for differences between hhMb and swMb. It is the third component ***U***_3_ that clearly distinguishes the XFEL datasets from the synchrotron datasets in a temperature-dependent manner (Fig. 4ace). In other words, the cryo structures (1dwr-t), ambient temperature structures (1bz6/p/r) including the deoxy hhMb at 7°C (5d5r), and the room temperature XFEL structures line up in an ascending order along the dimension *c*_3_. Not incidentally, the photolyzed structures Mb*CO and **CO have the most negative coefficient *c*_3_ because they were obtained at temperatures even lower than 100 K (Chu et al., 2000). The corresponding decomposed map ***U***_3_ reveals the thermal effects in electron densities of the heme as a nearly symmetric shell of spherical positive densities around the iron with very little positive or negative density on the iron itself (Fig. 6), which transitions into a square shape with four corners centered at the six-membered rings in the heme plane (marked 1-4 in Fig. 6e). This symmetric ***U***_3_ of the omit maps shows no usual signals for ligand dissociation and heme doming therefore contrasts with the asymmetric ***U***_4_ of the difference maps characterized by the distinct cube-shaped positive density maxima at the eight corners. Two sheets of positive densities are found at the cross sections parallel to, and 1.3 Å away from, the heme plane on both sides (Fig. 6df). They are connected via pillar-shaped positive densities that run through the six-membered rings (marked 1-4 in Fig. 6e) and the pyrrole rings (marked 5-8 in Fig. 6e) along with some other positive densities outside the heme (marked 9-20 in Fig. 6e). It is noteworthy that such thermal effects do not stand out as a major component in the SVD analysis of difference maps because they are largely self-canceled during the subtraction of the reference.

**Figure 6.**
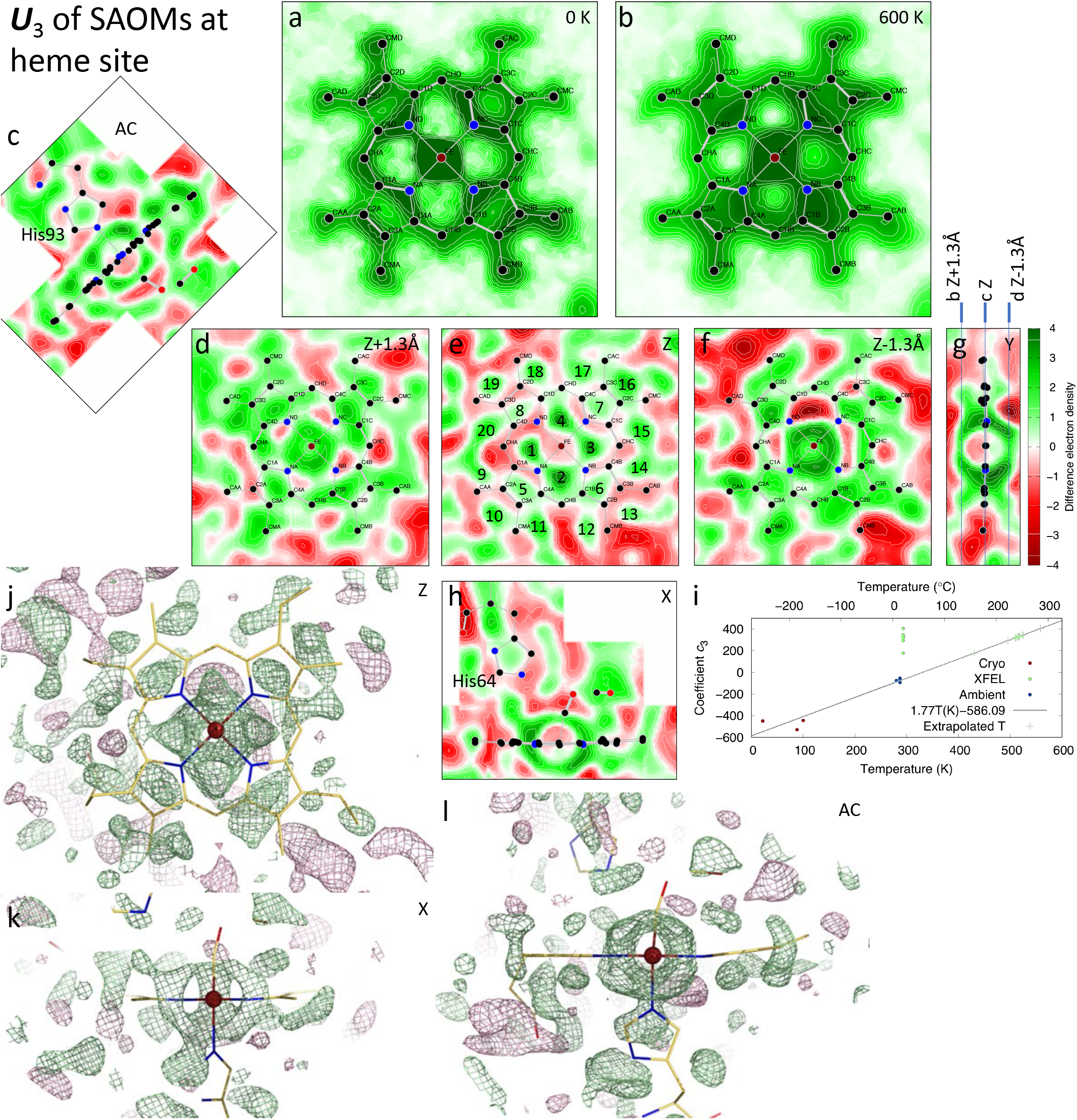
Thermal vibrations isolated from the chemical signals of photolysis. The third component of the SAOMs ***U***_3_ is highly correlated with the sample temperature (Fig. 4ace). A linear extrapolation based on the datasets available at cryo and ambient temperatures predicts that the instantaneous local temperature jump of the heme may have exceeded 500 K in the XFEL datasets (i). (a and b) Linear combinations of 900***U***_1_-586***U***_3_ and 900***U***_1_ + 476***U***_3_ reconstitute the omit maps of the heme at 0 and 600 K, respectively, excluding all chemical signals related to photolysis, where the coefficient *c*_1_ = 900 of ***U***_1_ is nearly constant for all omit maps. The transition from 0 to 600 K free of the chemical signals from photolysis is shown in Movie S2. (c-h) Cross sections through the third component ***U***_3_. Positive densities in 20 pillars connecting two sheets of positive densities above and below the heme in the cross sections of *Z*±1.3 Å are marked in (e). (j-l) Three-dimensional meshes of this component in green and red are contoured at ±15*σ*, respectively. See Fig. 3 legend for more detail.

The temperature dependency and the symmetry of the signals in ***U***_3_ strongly suggest that these features are related to the local temperature at which the MbCO molecules were probed, rather than artifacts arising from structural refinement or difference in spatial resolutions among datasets. All datasets used for the analysis are unified to a similar spatial resolution (Methods). The positional parameters of the models within the omitted region and the thermal parameters of the entire globin are intentionally “forgotten” in the calculation of the omit maps. The large *c*_3_ coefficients of the XFEL datasets are due to instantaneous temperature elevation at the heme site upon exposure to each ultrashort XFEL pulse, which introduces isotropic vibrations at high frequencies to the entire tetrapyrrole cofactor as captured by ***U***_3_. This molecular image of ***U***_3_ shows an increasing intensity from cryo to ambient temperatures and to a very high local temperature induced by XFEL pulses (Movie S2). Solely based on an estimate by linear extrapolation without other validations, the instantaneous temperature of the heme may exceed 500 K during an 80-fs XFEL pulse (Fig. 6i). However, it remains to be seen whether this analysis of the omit maps restricted within the heme site can be extended to show that the temperature of the globin or the crystals has a significant elevation during an XFEL pulse.

Identification of such symmetric and temperature-dependent signals is important for interpretation of the highly dynamic ultrafast signals. The fact that the symmetric ***U***_3_ and the asymmetric ***U***_7_ of the omit maps are orthogonal to each other mathematically guarantees that these coexisting components are not interchangeable and not cross contaminated between them (Methods). The cubic shaped ***U***_7_ of the omit maps and its counterpart ***U***_4_ of the difference maps are the chemical consequence of ligand dissociation; the symmetrical and spherical shaped ***U***_3_ originates from the thermal consequence. One cannot compensate for the other and both coexist in all maps. The equivalents can be stated: The cubic positive densities influenced by the 3*d* orbitals are not due to thermal vibrations; the symmetrically enlarged iron and the thickened heme due to an instantaneous temperature elevation have been isolated from the chemically relevant signals. It is worth noting that the ultrafast onset of ***U***_4_ of the difference maps (Fig. 2cd) is not in contradiction with ***U***_7_ of the omit maps contained not only in time-resolved XFEL datasets but also in the static structures of deoxy Mb and photolyzed Mb*CO and **CO at different temperatures (Fig. 4e). Instead, regaining of the high-spin 3*d* orbitals of the iron occurs before the first time point at tens of fs; this permanent change lasts throughout the dissociation reaction into the deoxy product.

The asymmetric ***U***_7_ of the omit maps also describes an important but unfamiliar structural event upon the ligand dissociation – the heme in-plane movement. This type of motion is harder to capture in difference maps due to little net gain or loss of electron density that would be caused by an in-plane movement of the heme. With respect to the protein framework, the heme moves towards its AB rings in addition to the other changes (Movie S3). Heme in-plane sliding is consistent with the strong signals of the skewed iron displacement and the proximal His93 (Fig. S3) that are closely related to structural responses in the globin. Similar in-plane sliding is even more significant in tetrameric hemoglobins (Baldwin and Chothia, 1979), which is the origin of the cooperative oxygen affinity (Ren, 2013b). However, the functional role of the heme in-plane sliding in Mb is not yet clear and this discussion branches away from the topic here on ultrafast structural changes.

### Global oscillation at low frequency

A recent solution scattering experiment using XFEL pulses showed that the radius of gyration *R*g first increases within 1 ps upon photodissociation of MbCO and then oscillates with a period of 3.6 ps, equivalent to 9 cm^-1^, a frequency lower than those observed by coherence spectroscopy (Levantino et al., 2015b). The molecular volume increases afterward with a phase shift. These observations indicate a wavelike motion propagating outward in the protein. Recent molecular dynamics simulations have confirmed that a pressure wave generated by photolysis reaches the molecular surface at 300 fs along the direction perpendicular to the heme plane (Brinkmann and Hub, 2016), which corresponds to a frequency of 28 cm^-1^ if the wave is reflected by the surface and oscillates. It was found by the simulations that the oscillation in solvent would be overdamped thus dissipates quickly.

Some SVD components derived here display strict correlation suggesting their dependency on each other. The ***U***_2_-***U***_3_ pair of the difference maps (Fig. 2b) is evidently equivalent to the ***U***_5_-***U***_8_ pair of the omit maps (Fig. 4b). When the difference maps of the entire globin are analyzed, the same signals emerge as the top-ranked ***U***_1_-***U***_3_ pair suggesting that the local signals at the heme site concur with global signals over the entire globin (Fig. 7). Interestingly, this pair of components displays a near-perfect circular correlation at sub-ps delays but is almost absent at longer delays > 1 ps (Fig. 7afgh). Such an oscillation at a low frequency requires at least two orthogonal components to depict. The oscillating signals have spread over the entire globin at sub-ps delays while their amplitudes decay to nearly zero after a few ps. Based on least-squares fittings of the corresponding coefficients (Methods), the frequency of the oscillation is determined to be 36±1 cm^-1^, corresponding to a period of 0.93 ps. The equivalent oscillation frequencies determined from the difference maps and omit maps at the heme site are 42±2 cm^-1^ (a period of 0.8 ps) and 37±1 cm^-1^ (a period of 0.89 ps), respectively.

**Figure 7.**
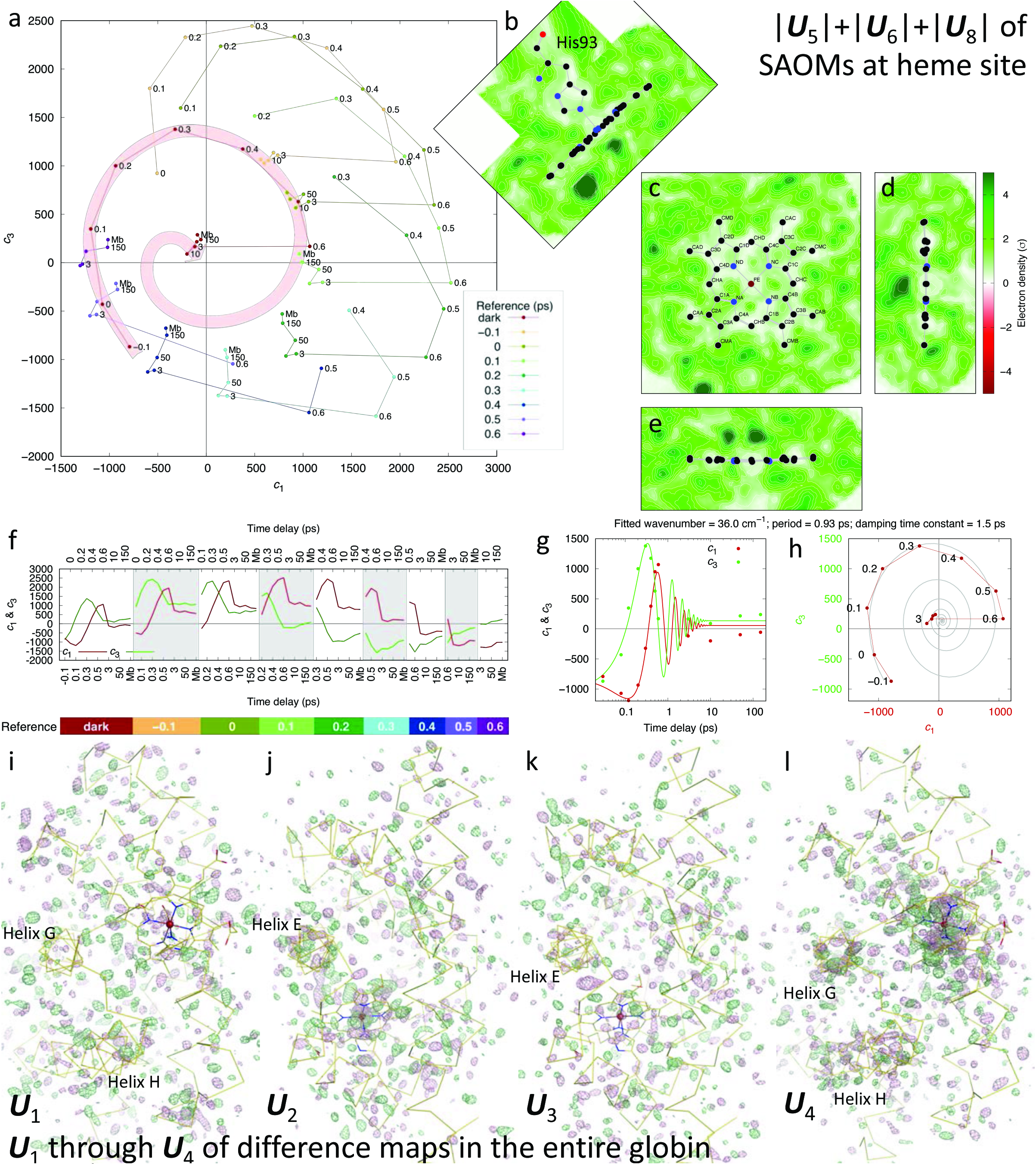
Low frequency oscillations in the solvent. (a) SVD analysis of the difference maps in the entire globin produces the first and third components circularly oscillating before 1 ps and quickly diminishing during the first few ps. Similar oscillations are also observed only around the heme site in both the difference maps (Fig. 2b) and the omit maps (Fig. 4b). (b-e) Cross sections through an averaged map of both the positive and negative halves of ***U***_5_, ***U***_6_, and ***U***_8_ of the SAOMs. Low densities in white and light green are located consistently on the heme and protein, which suggests that the oscillating signals are more associated with the solvent rather than the globin. (f) The circularly oscillating coefficients *c*_1_ and *c*_3_ are plotted as function of delay time in many series. The offset sinusoidal curves indicate a circular oscillation. Each map series is derived from subtracting a different reference dataset as indicated by the color bar below. (g and h) *c*_1_ and *c*_3_ of the -*F*_dark_ series are least-squares fitted with a two-dimensional damped oscillation (Methods). (i-l) Comparison of several components of difference maps in the entire globin. ***U***_1_ and ***U***_3_ contain widespread signals, as they are ranked the most significant components. But no signal is concentrated on the heme or the protein. Instead, ***U***_2_ and ***U***_4_ clearly contain strong signals associated with the heme and secondary structures of the protein.

The low frequency oscillation determined from the XFEL datasets of Barends & Schlichting et al. is highly comparable to those measured by fs vibrational coherence spectroscopy (Gruia et al., 2008; Rosca et al., 2002). I postulate that this low frequency reflects the molecular size of the solvated Mb rather than the property of the heme. First, the corresponding period of 0.93 ps is about the time that a mechanical wave takes to traverse 14 Å, that is, an average distance from the heme to the molecular surface of Mb, given a speed of sound at 1.5 km/s or nm/ps (Fig. S1). Second, the molecular sizes of several heme proteins, such as cytochrome c (12 kDa), Mb (17 kDa), and cystathionine β-synthase (63 kDa), show good anticorrelation with their dominant low frequencies of 44, 40, and 25 cm^-1^, respectively (Gruia et al., 2008; Karunakaran et al., 2010, 2014). Third, no association between the oscillating signals and specific structural features can be found, such as the heme, ligand, surrounding side chains, or helices (Fig. 7ik). These oscillating signals sharply contrast with strong, non-oscillating signals associated with the heme and various helices evidenced by the second and fourth components of the difference maps (Fig. 7jl), where helices E, G, and H move away from the heme in response to ligand dissociation (Barends et al., 2015).

These motion signals of various structural elements are much enhanced by SVD. The oscillating signals at low frequency seem to propagate via solvent within the protein matrix rather than the globin. Strong positive and negative peaks in ***U***_5_, ***U***_6_, and ***U***_8_ of the omit maps are consistently found in the solvent region away from the heme and protein. However, a specific description on how the solvent responses to ligand dissociation is difficult, since the solvent structure is far less well determined than the globin structure. An overall contribution of ***U***_5_, ***U***_6_, and ***U***_8_ can be judged by a sum of the absolute values |***U***_5_|+|***U***_6_|+|***U***_8_| that enhances the distribution of signals. The heme and globin are located in low intensities of these oscillating components (Fig. 7b-e). These global oscillations occur only a few times before dissipation (Gruia et al., 2008; Rosca et al., 2002). However, there are not enough time points to determine the damping time constant, which is roughly estimated to be 1-4 ps. Contrary to the common belief that low frequency oscillations are due to the properties of the heme, here I attribute the low frequency oscillation to the global response of protein and solvent as a whole to a sudden molecular event – photodissociation of the ligand, where the vibrational energy released from the photodissociation is quickly dissipated into the environment.

## Discussion

### Are atomic orbitals observed?

It requires ultrahigh spatial resolution in a Fourier synthesis to accurately depict the probability distribution of a single electron occupying its orbital. For example, iron 3*d* and sulfur 3*p* densities in an Fe4S4 cluster are described with crystallographic data at a resolution of 0.48 Å with an analytical technique called charge-density analysis (Hirano et al., 2016), in which atoms are considered as aspherical and modeled by spherical harmonic functions in contrast to spherical Gaussian functions used in everyday protein crystallography (Coppens, 1998). Since the non-bonding inner electrons remain largely constant in a spherical distribution as a whole, the inner electrons in a majority can easily overwhelm the presence of the bonding electrons in a minority, let alone subtle changes in electron transfer. Therefore, it is very difficult, if not impossible, to detect small orbital changes upon bond rupture using regular electron density maps such as a 2Fo-Fc map. Difference Fourier technique is required as it is far more sensitive to small changes (Henderson and Moffat, 1971), in which the constant, spherical distribution of the inner electrons is self-canceled and the remaining densities would reveal electron transfer. However, the reality is usually far less ideal because of conformational changes. Atomic displacements involved in a conformational change carry all electrons associated with the moving atoms and result in significant positive and negative densities in a difference Fourier map, that is, a spatial distribution of any gain and loss of electrons. Atomic displacements such as a side chain rotation or the heme iron movement by a few tenths of an Å again conceal the subtle signal of an electron transfer from one orbital to another.

The difference maps derived here as SVD components represent partial, variable signals in addition to the constant electron density map regardless of the bonding change. An ultrahigh spatial resolution is required to depict slightly aspherical features when an overwhelming spherical background is present. Here, a combined use of difference Fourier technique and decomposition eases the demand for spatial resolution to reveal subtle changes. Furthermore, the notion that an ultrahigh spatial resolution is required for observations of orbital changes contradicts the fact that there is no discontinuity in the probability distribution of any electronic orbital. A reasonably high spatial resolution shall be sufficient to analytically describe the smooth function of a gain or loss of an electron. Therefore, the small amplitude of desired changes over a much greater constant background hinders a reliable observation more than the lack of high spatial frequencies does. This work presents the effectiveness of the strategy that enables numerical deconvolution to resolve concurrent events so that the image of a partial change, such as a single electron transfer, can be isolated from other changes. At an available spatial resolution, the partial change due to a specific electronic event deconvoluted from much greater conformational changes could be the remanece of orbital densities instead of an accurate image of these orbitals.

A limited experimental spatial resolution also raises a doubt whether some unusual features are the artifacts due to a Fourier truncation error. A Fourier truncation error may occur to several static datasets included in this analysis, in which the spatial resolutions are truncated to 1.75 Å to match that of XFEL datasets. Such errors may appear as minor components after SVD (Figs. 2e and 4f). When the average intensity of an experimental dataset naturally decays as function of the spatial resolution, Fourier truncation errors are damped to a minimum. Difference Fourier technique again self-cancels the remaining errors because both datasets involved in the subtraction would carry similar Fourier truncation errors if any. In addition, the applied weighting scheme (Ren et al., 2001, 2013; Šrajer et al., 2001; Ursby and Bourgeois, 1997) severely reduces any large differences in Fourier synthesis (Methods), which also prevents Fourier truncation errors. After the decomposition, proper rotations in the SVD space validate that the signals under discussion, such as the positive cubic network, is not due to Fourier truncation errors because of its correlation with the metadata. An artifact due to Fourier truncation errors, if so conclude, needs to be explained why it is correlated with a physical or chemical parameter.

The theoretical radii of the 3*d* orbitals for an isolated Fe(II) cation are about 0.36 Å at the maximum probability of these orbitals. However, the orbital densities spread out many folds beyond the maxima (Waber and Cromer, 1965). The majority of the iron 3*d* orbital densities in an Fe4S4 cluster span about 1 Å from the nucleus as measured in a protein (Hirano et al., 2016). The observed distances between the iron and the corners of the observed cubic densities are significantly larger. Therefore, the observed cubic densities here are not an accurate image of a probability distribution of the 3*d* electrons. However, this density distribution, given the available spatial resolution, is clearly influenced by the unpaired 3*d* electrons. The net gain of electron densities could be further away from the nucleus compared to the maximum probability of these orbital. For a practical purpose in protein crystallography, an accurate depiction of a 3*d* orbital is not the task. Rather, direct observations of a specific electronic event would provide unprecedented insight into the essence of biochemistry.

### Temporal resolution vs. resolution of heterogeneity

Ultrashort XFEL pulses improve the time resolution to sub-ps or even tens of fs compared to 100 ps at synchrotrons. This ultrafast time resolution, when combined with the spatial resolution of crystallography, offers an unprecedented opportunity for direct observation of transient structural changes in a photochemical reaction as detail as individual electrons. However, due to their highly dynamic and heterogeneous nature, it is difficult to experimentally isolate short-lived structural species along the temporal dimension regardless of the ultrafast time resolution. In other words, resolving the population difference between two chemical species at one fs from the next does not mean that one can resolve two distinct structures. As demonstrated by this work, analytical methods such as SVD are needed to deconvolute them from mixtures of intermediate species. The strategy we proposed for resolution of structural heterogeneity (Ren et al., 2013) is highly effective for separating independent electronic and atomic events that often coexist during a chemical reaction or a biological process.

This work shows that it is possible to observe structural changes as small as a gain or loss of individual electrons by protein crystallography. Small structural signals can be correlated with their corresponding metadata by proper rotations, that is, deconvolution after decomposition, thus allowing the origin of these signals be revealed. I posit that all observations so far are the consequence, rather than the cause, of the ligand dissociation. The proposed transient event of metal-to-ligand charge transfer is not captured here, presumably because the transferred electron is delocalized all over the porphyrin (Franzen et al., 2001). Therefore, the fundamental questions remain for photolysis of MbCO: Upon the absorption of a quantum of energy from a green photon, what initial electronic events lead to a partitioning of the bonding electrons in two separate atoms, that is, the bond rupture? What electronic events cause spin crossover? To image these events directly, even higher time resolution is required to capture electron density changes prior to and during bond rupture.

### Future outlook of ultrafast dynamic crystallography

Despite the recent developments in SFX and its applications to time-resolved studies of a few model systems, it is yet premature to declare that the current SFX platform applies to other systems of more significant biological importance such as visual and microbial rhodopsins, plant and bacterial phytochromes, photosynthetic reaction centers, and light-dependent DNA repair enzymes. There are at least three major challenges. First, it is not practical to produce and inject an astronomical number of nano or microcrystals of these proteins into an XFEL beam to support data collections of a time series. The current serial protocol used to obtain the MbCO and other data has a diminishingly small yield of useful diffraction data, based on the ratio between the number of diffracting crystals and the total number of crystals produced (Ren et al., 2018). That is to say, the vast majority of the crystals have no chance to diffract X-ray even when they are injected into the X-ray beam. Such a yield, defined differently from the hit rate, is not sustainable for other photosensitive protein crystals. Several “fixed target” implementations of crystal delivery systems have been reported, including one that we have recently proposed (Ren et al., 2018). However, it remains to be seen how effective they are in achieving a higher yield while maintaining the diffraction quality.

Second, since the XFEL pulses usually carry insufficient bandwidths to fully integrate the Bragg reflections and in the meanwhile angular rotation of a crystal during an ultrashort XFEL pulse is not attainable, tens to hundreds of thousands of diffraction images are required to obtain integrated intensities for all Bragg reflections in a complete sampling of the reciprocal space. A possibility to solve this partiality problem is to use an X-ray beam with a significant convergence of about a degree (Ho et al., 2002; Spence et al., 2014), which will produce completely integrated intensities with a far fewer number of diffraction images, thereby alleviating the demand for a large quantity of purified proteins.

Third, data merging from a large crystal pool is a major source of error in producing difference signals for photosensitive crystals, particularly those with much lower diffraction power than Mb crystals. To address this challenge, we also proposed a protocol to obtain difference signals that bypasses the integrated intensities of Bragg reflections. We reason that integrated intensities are not necessary if the dark and light diffraction images can be obtained within a single exposure of an XFEL pulse because the percentage change in integrated intensities of a given reflection, that is, the time-resolved signal, is already captured even if this reflection is partially measured in both dark and light conditions. A plan has been laid to implement this protocol, based on a dichromator instrument that achieves angular split and temporal delay at the same time (Ren and Yang, 2016). When each of these technical hurdles is overcome, direct observations of atomic and electronic movements during ultrafast processes will be closer to broad applications for many light sensitive crystals of great biological and medical significance.

## Code availability

Computer software dynamiX™ is available from Renz Research, Inc.

## Acknowledgements

The following database and software are used in this work: CCP4 (ccp4.ac.uk), Coot (www2.mrc-lmb.cam.ac.uk/Personal/pemsley/coot), dynamiX™ (Renz Research, Inc.), gnuplot (gnuplot.info), Mathematica (wolfram.com), PDB (rcsb.org), PHENIX (phenix-online.org), PyMOL (pymol.org), Python (python.org), and SciPy (scipy.org).

## Methods

### Difference Fourier maps

Difference Fourier maps are commonly used in protein crystallography. A difference Fourier map is synthesized from a Fourier coefficient set of *F*_light_-*F*_reference_ with the best available phase set. Before Fourier synthesis, *F*_light_ and *F*_reference_ must be properly scaled to the same level so that the distribution of difference values is centered at zero and not skewed either way. All available datasets from Barends & Schlichting et al. (2015) are carefully scaled and difference Fourier maps are calculated using a software package dynamiX™ according to previously published protocols including a weighting scheme under the assumption that a greater amplitude of a difference Fourier coefficient *F*_light_-*F*_reference_ is more likely caused by noise than by signal (Ren et al., 2001, 2013; Šrajer et al., 2001; Ursby and Bourgeois, 1997).

Both the dark and light datasets can serve as a reference in difference maps. If a light dataset at a certain delay is chosen as a reference, the difference maps show the changes since that delay time but not the changes prior to that delay. These 91 maps in a full matrix provide independent observations for all changes (Table S1). This strategy is important to the subsequent singular value decomposition (SVD) analysis. In addition, two more difference maps of the photolyzed Mb*CO and **CO (1dws/t) are also included in the -*F*_dark_ series for SVD analysis.

Each difference map is masked around the heme site. All densities at those grid points are set to zero, if they are greater than 5 Å away from the heme, the bound or docked CO, the side chain of the proximal His93, and that of the distal His64. Difference maps around the heme site include the most significant signals and exclude noises in the protein and solvent. However, the unmasked difference maps in the entire globin are also used a separate SVD analysis to include the protein and solvent regions for observation of global changes (Fig. 7i-l).

### Simulated annealing omit maps

The heme, the ligand, the side chains of proximal His93 and distal His64 are removed from all structures listed in Table S2. The remaining structures are refined using simulated annealing method with phenix.refine starting from a simulated temperature of 5,000 K. Fourier coefficients are obtained from the differences between the structure factor amplitudes *F*_observed_ observed in the time-resolved experiments and the structure factor amplitudes *F*_calculated_ calculated from the omitted structures. The resulting *F*_observed_-*F*_calculated_ maps, known as simulated annealing omit maps (SAOMs), reveal the unbiased electron densities in the omitted region. Since the sperm whale (sw) Mb structures are in a different unit cell compared to the horse heart (hh) Mb structures, the SAOMs are aligned together according to a least-squares fitting of the main chain atoms in residues 2-151. The heme and ligand are not used in the alignment procedure. Therefore, the motion of heme found in these omit maps is with respect to the globin framework (Fig. 5 and Movie S3). 20 aligned omit maps are subjected to the subsequent SVD analysis.

### Singular value decomposition of (difference) electron density maps

The principle of SVD can be found in texts on linear algebra. SVD analysis of (difference) electron density maps was previously described (Ren et al., 2013; Schmidt et al., 2003). Here I briefly recap this numerical procedure with an emphasis on the physical meaning of the outcome.

An electron density map, sometimes a difference map, consists of density values on a collection of grid points within a mask (see above), also known as voxels that are equivalent to pixels in a digitized image. All *M* voxels in a three-dimensional map can be serialized into a one-dimensional sequence of density values according to a specific protocol. It is not important what the protocol is as long as a consistent protocol is used to serialize all maps of the same grid setting and size, and a reverse protocol is available to erect a three-dimensional map from a sequence of *M* densities. Therefore, a set of *N* serialized maps, also known as vectors in linear algebra, can fill the columns of a data matrix **A** with no specific order so that the width of **A** is *N* columns and the length is *M* rows. Often, *M* >> *N*, thus **A** is an elongated matrix. In this work, *N* = 93 for the difference maps and *N* = 20 for the omit maps. If a consistent protocol of serialization is used, the corresponding voxel in all *N* maps occupies a single row of matrix **A**. For example, the voxels at the heme iron in all maps could occupy the *j*th row of matrix **A**. This strict correspondence in a row of matrix **A** is important. Changes of the density values in a row are due to either signals, systematic errors, or noises.

SVD of the data matrix **A** results in **A** = **UWV^T^** also known as matrix factorization. Matrix **U** has the same shape as **A**, that is, *N* columns and *M* rows. The *N* columns contain decomposed basis components ***U****_k_*, known as left singular vectors of length *M*, where *k* = 1, 2, …, *N*. Therefore, each component ***U****_k_* can be erected using the reverse protocol to form a three-dimensional map. This decomposed elemental map can be presented in the same way as the original maps, for example, rendered in molecular graphics software such as Coot and PyMol. The second matrix **W** is a square matrix that contains all zeros except for *N* positive values on its major diagonal, known as singular values *w*_*k*_. The magnitude of *w*_*k*_ is considered as a weight or significance of its corresponding component ***U****_k_*. The third matrix **V** is also a square matrix of *N* × *N*. Each column of **V** or row of its transpose **V^T^**, known as a right singular vector, contains the relative compositions of ***U****_k_*. That is to say, the *N* values in a column ***V***_*k*_ describe the relative compositions of the component ***U****_k_* in the *N* original maps. A singular triplet denotes 1) a decomposed component ***U****_k_*, 2) its singular value *w*_*k*_, and 3) the composition function ***V***_*k*_. Singular triplets are often sorted in a descending order of their singular values *w*_*k*_. Only a small number of *n* significant singular triplets identified by the greatest singular values *w*_1_ through *w_n_* can be used in a linear combination to reconstitute a set of composite maps that closely resemble the original ones in matrix **A**, where *n* < *N*. For example, the original map in the *i*th column of matrix **A** can be closely represented by the *i*th composite map *w*_1_*v*_1*i*_***U***_1_ + *w*_2_*v*_2*i*_***U***_2_ + … + *w_n_v_ni_**U**_n_*, where (*v*_1*i*_, *v*_2*i*_, …) is the *i*th row of matrix **V**. *c_k_* = *w_k_v_ki_* in front of ***U****_k_* is the coefficients for the linear combination. Excluding the components after ***U***_*n*_ in this linear combination is based on an assumption that the singular values after *w_n_* are very small relative to those from *w*_1_ through *w_n_*. As a result, the structural information evenly distributed in all *N* original maps is effectively concentrated into a far fewer number of *n* significant components, known as information concentration or dimension reduction. In this study, the difference maps seem to contain 15 major components; the top nine are the most significant (Fig. 2e). The components 10 through 15 may very likely harbor less and less signals or systematic differences. For example, Fourier truncation errors, if significant enough, could occupy one or more of these components. However, the components 10 through 15 have not been examined in detail.

On the other hand, the trailing components in matrix **U** contain inconsistent fluctuations and random noises. Excluding these components effectively rejects noises. However, no clear boundary is guaranteed between signals, systematic errors, and noises. Systematic errors could be more significant than the desired signals, for example, the second and fourth components ***U***_2_ and ***U***_4_ of the omit maps describe significant differences between hhMb and swMb, a valid scientific point to investigate but outside the scope of this paper.

### The orthonormal property of SVD

The solution set of SVD must guarantee that the columns in **U** and **V**, the left and right singular vectors ***U****_k_* and ***V***_*k*_, are orthonormal, that is, ***U****_h_*·***U****_k_* = ***V****_h_*·***V***_*k*_ = 0 (ortho) and ***U****_k_*·***U****_k_* = ***V***_*k*_·***V***_*k*_ = 1 (normal), where *h* ≠ _*k*_ but both are from 1 to *N*. The orthonormal property also holds for the row vectors. As a result, each component ***U****_k_* is independent of the other components. In other words, a component cannot be represented by a linear combination of any other components. For example, the symmetrical, spherical, and thermal component ***U***_3_ (Fig. 6) and the asymmetrical, cubic component of ***U***_7_ (Fig. 5) of the omit maps are orthonormal. Both components show positive densities around the iron. But the cubic positive densities in ***U***_7_ are associated with the strong negative densities on the ligand depicting dissociation, which is a direct link from ligand dissociation to regaining of the high-spin 3*d* orbitals of the iron. On the other hand, no sign of ligand dissociation is found in the component ***U***_3_ that features the spherical densities, which strongly suggests that this component describes the thermal effect. These two components are not interchangeable despite some similarity in the arrangement of positive densities, that is, one of the physical meanings of the orthonormal property of SVD. However, two physical or chemical parameters, such as temperature and pH, may cause different changes to a structure, but not necessarily orthogonal changes (see below).

Due to the orthonormal property of SVD, an *N*-dimensional Euclidean space is established, and the first *n* dimensions define its most significant subspace. Each coefficient set *c_k_* = *w_k_v_ki_* of the *i*th composite map is located in this subspace. All coefficient sets for *i* = 1, 2, …, *N* in different linear combinations to approximate the *N* original maps in a least-squares sense can be represented by *N* points in the *n*-dimensional Euclidean subspace. In this work, many two-dimensional orthographical projections of this *n*-dimensional subspace are presented as scatter plots with each map represented as a dot located at a position determined by the coefficient set of this map. These scatter plots are highly informative to reveal the relationship between the (difference) electron density maps and their metadata. For example, the difference maps at short time delays < 1 ps are consistently different from those at long time delays > 1 ps in a certain aspect (Fig. 2ac) and that two major components are circularly correlated but only before 1 ps, that is, a damped oscillation (Fig. 2b).

### Rotation in SVD space

Dimension reduction is indeed effective in data analysis of dynamic crystallography when many datasets are evaluated at the same time. However, the interpretation of a basis component ***U****_k_*, that is, “what-does-it-mean”, requires a clear demonstration of the relationship between the core data, here (difference) electron density maps, and their metadata, such as time, temperature, and other experimental conditions. The outcome of SVD does not guarantee any physical meaning in a basis component. Therefore, SVD provides no direct answer to “what-does-it-mean”. The factorized set of matrices **U**, **W**, and **V** from SVD is not a unique solution. That is to say, they are not the only solution to factorize matrix **A**. The default solution set from SVD may not carry an obvious physical meaning to depict “what-does-it-mean”, thus its usefulness is very limited to merely a mathematical construction. A more straightforward analogy is that one can walk along two perpendicular streets to reach the opposite corner of a street block as if walking a diagonal block despite the fact that a diagonal street does not physically exist. This is also the origin of the debate on whether atomic orbitals as a basis set of mathematical constructions are physical and observable (Scerri, 2000). Therefore, it is very important to find one or more alternative solution sets that are physically meaningful to elucidate a structural interpretation. The concept of rotation after SVD was introduced by Henry & Hofrichter (1992). But they suggested a protocol that fails to preserve the orthonormal and least-squares properties of SVD. The rotation process, also known as Givens rotation, is a connection between the analytical tool SVD and scientific findings. This rotation shall not be confused with a rotation in the three-dimensional real space, in which a molecular structure resides.

A rotation in the *n*-dimensional Euclidean subspace is necessary to change the perspective before a clear relationship emerges to elucidate scientific findings. It can be shown that two linear combinations are identical before and after a rotation of an angle *θ* in a two-dimensional subspace of *h* and *k*. That is,

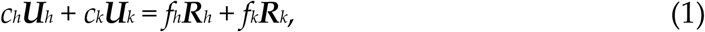

where *c_h_* and *c_k_* are the coefficients of the components ***U***_*h*_ and ***U***_*k*_ before the rotation as defined above; and *f_h_* and *f_k_* are the coefficients of the rotated basis components ***R****_h_* and ***R****_k_*, respectively. The same Givens rotation is applied to both the bases and their coefficients:

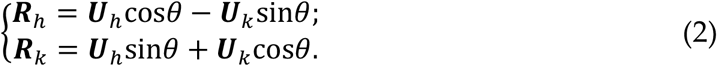

Obviously, the rotated components ***R****_h_* and ***R****_k_* remain mutually orthonormal and orthonormal to other components. And

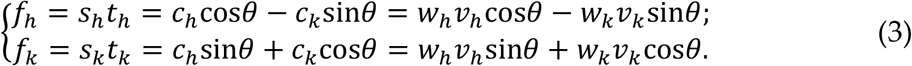

Here 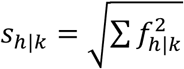 are the singular values that replace *w_h_* and *w_k_*, respectively, after the rotation. They may increase or decrease compared to the original singular values so that the descending order of the singular values no longer holds. ***T***_*h*|*k*_ = (*t*_*h*|*k*1_, *t*_*h*|*k*2_, …, *t*_*h*|*kN*_) = (*f*_*h*|*k*1_, *f*_*h*|*k*2_, …, *f*_*h*|*kN*_)/*s*_*h*|*k*_ are the right singular vectors that replace ***V***_*h*_ and ***V***_*k*_, respectively. ***T***_*h*_ and ***T***_*k*_ remain mutually orthonormal after the rotation and orthonormal to other right singular vectors that are not involved in the rotation.

A rotation in two-dimensional subspace of *h* and *k* has no effect in other dimensions, as the orthonormal property of SVD guarantees. Multiple Givens rotations can be carried out in many two-dimensional subspaces consecutively to achieve a multi-dimensional rotation. Rotation in the SVD space converts one solution set **A** = **UWV^T^** to other alternative solutions **A** = **RST^T^** so that an appropriate perspective can be found to elucidate the relationship between the core data and metadata clearly and concisely. A solution derived from a rotation retains the orthonormal property of SVD. SVD analyses presented in this paper employ rotations extensively except that no distinction is made in the symbols of components and coefficients before and after a rotation.

For example, a large gap between all difference maps < 1 ps and those > 1 ps indicates major differences between these two distinct time scales (Fig. 2a). A rotation does not change this fact except that this large gap used to orient along some inclined direction before the rotation. Therefore, the original component ***U***_1_ cannot describe the structural differences between the short delays < 1 ps and the longer delays > 1 ps, because the original ***U***_1_ used to be a linear combination of several factors. The best rotation is to separate these two groups of difference maps along a major dimension, that is, a deterministic solution. The large gap exists as soon as an SVD analysis is performed and can be easily revealed in an interactive three-dimensional plotting on the computer screen, although it is not obvious in any of the orthographical projections before a proper rotation is found. One cannot create nor eliminate this large gap by any rotation. A different rotation may reorient this large gap along another major dimension. But the structural conclusion would be equivalent.

All difference maps have their coefficients *c*_4_ ≈ 0 except those calculated with the dark dataset as a reference (red trace in Fig. 2cd). The best rotation can be deterministically obtained by fitting all difference maps into the plane of *c*_4_ = 0 except the -*F*_dark_ series standing outside the plane. The resulting component ***U***_4_ depicts features only in the -*F*_dark_ series (Fig. 3). Again, the fact cannot be changed with any rotation that light-dark difference maps standout from the other difference maps arranged on a plane that is not necessarily *c*_4_ = 0 before a proper rotation. Therefore, the original ***U***_4_ are hard to interpret with several physical meanings fixed in it. In this sense, it is the rotation after the decomposition that completes the deconvolution of physical and chemical causes. See also an example of rotation in an SVD analysis of atomic coordinates of protein structures (Ren, 2016).

### Least-squares fittings of oscillating components

Oscillating components are repeatedly found in the difference maps and omit maps. Usually, a pair of components are oscillating with the same frequency, which does not exclude the possibility that a third weaker component is also oscillating at the same frequency. A pair of coefficients in a two-dimensional damped oscillation of a period *p* is least-squares fitted as *x* and *y*:

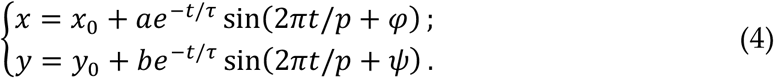

*x*_0_ and *y*_0_ are the center of the oscillation. *a* and *b* are the amplitudes of the elliptical oscillation in two dimensions. They are damped according to an exponential decay with a time constant *τ*. *φ* and *ψ* are the phase angles of the oscillation in two dimensions. They are not strictly *π*/2 apart.

The sixth and eighth components of the difference maps appear to gain a two-dimensional oscillation after 1 ps (Fig. 2f). Their coefficients are least-squares fitted as *x* and *y* with an exponential onset:

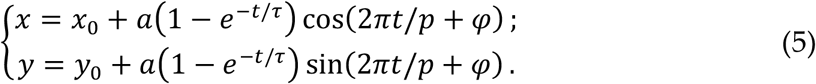

Due to the lack of sufficient time points, the oscillation is forced to take a circular form.

## Supplementary Table

**Table S1.** The full matrix of difference Fourier maps

| Reference (ps) | Dark | Time points (ps) |  |  |  |  |  |  |  |  |  |  |  |  |
| --- | --- | --- | --- | --- | --- | --- | --- | --- | --- | --- | --- | --- | --- | --- |
| <sup>1</sup>Dark (5cmv) | <sup>2</sup>. | -0.1 | 0 | 0.1 | 0.2 | 0.3 | 0.4 | 0.5 | 0.6 | 3 | 10 | 50 | 150 | deoxy |
| -0.1 (5cn4) | <sup>3</sup>- | . | 0 | 0.1 | 0.2 | 0.3 | 0.4 | 0.5 | 0.6 | 3 | 10 | 50 | 150 | deoxy |
| 0 (5cn5) | - | - | . | 0.1 | 0.2 | 0.3 | 0.4 | 0.5 | 0.6 | 3 | 10 | 50 | 150 | deoxy |
| 0.1 (5cn6) | - | - | - | . | 0.2 | 0.3 | 0.4 | 0.5 | 0.6 | 3 | 10 | 50 | 150 | deoxy |
| 0.2 (5cn7) | - | - | - | - | . | 0.3 | 0.4 | 0.5 | 0.6 | 3 | 10 | 50 | 150 | deoxy |
| 0.3 (5cn8) | - | - | - | - | - | . | 0.4 | 0.5 | 0.6 | 3 | 10 | 50 | 150 | deoxy |
| 0.4 (5cn9) | - | - | - | - | - | - | . | 0.5 | 0.6 | 3 | 10 | 50 | 150 | deoxy |
| 0.5 (5cnb) | - | - | - | - | - | - | - | . | 0.6 | 3 | 10 | 50 | 150 | deoxy |
| 0.6 (5cnc) | - | - | - | - | - | - | - | - | . | 3 | 10 | 50 | 150 | deoxy |
| 3 (5cnd) | - | - | - | - | - | - | - | - | - | . | 10 | 50 | 150 | deoxy |
| 10 (5cne) | - | - | - | - | - | - | - | - | - | - | . | 50 | 150 | deoxy |
| 50 (5cnf) | - | - | - | - | - | - | - | - | - | - | - | . | 150 | deoxy |
| 150 (5cng) | - | - | - | - | - | - | - | - | - | - | - | - | . | deoxy |
| Deoxy (5d5r) | - | - | - | - | - | - | - | - | - | - | - | - | - | . |
| PDB entry | 5cmv | 5cn4 | 5cn5 | 5cn6 | 5cn7 | 5cn8 | 5cn9 | 5cnb | 5cnc | 5cnd | 5cne | 5cnf | 5cng | 5d5r |

<sup>1</sup>Traditionally, the dark dataset is chosen as the reference in calculation of difference Fourier maps.
<sup>2</sup>Dot (.) indicates that these are empty maps.
<sup>3</sup>Dash (-) indicates that these are negated maps.

**Table S2.**
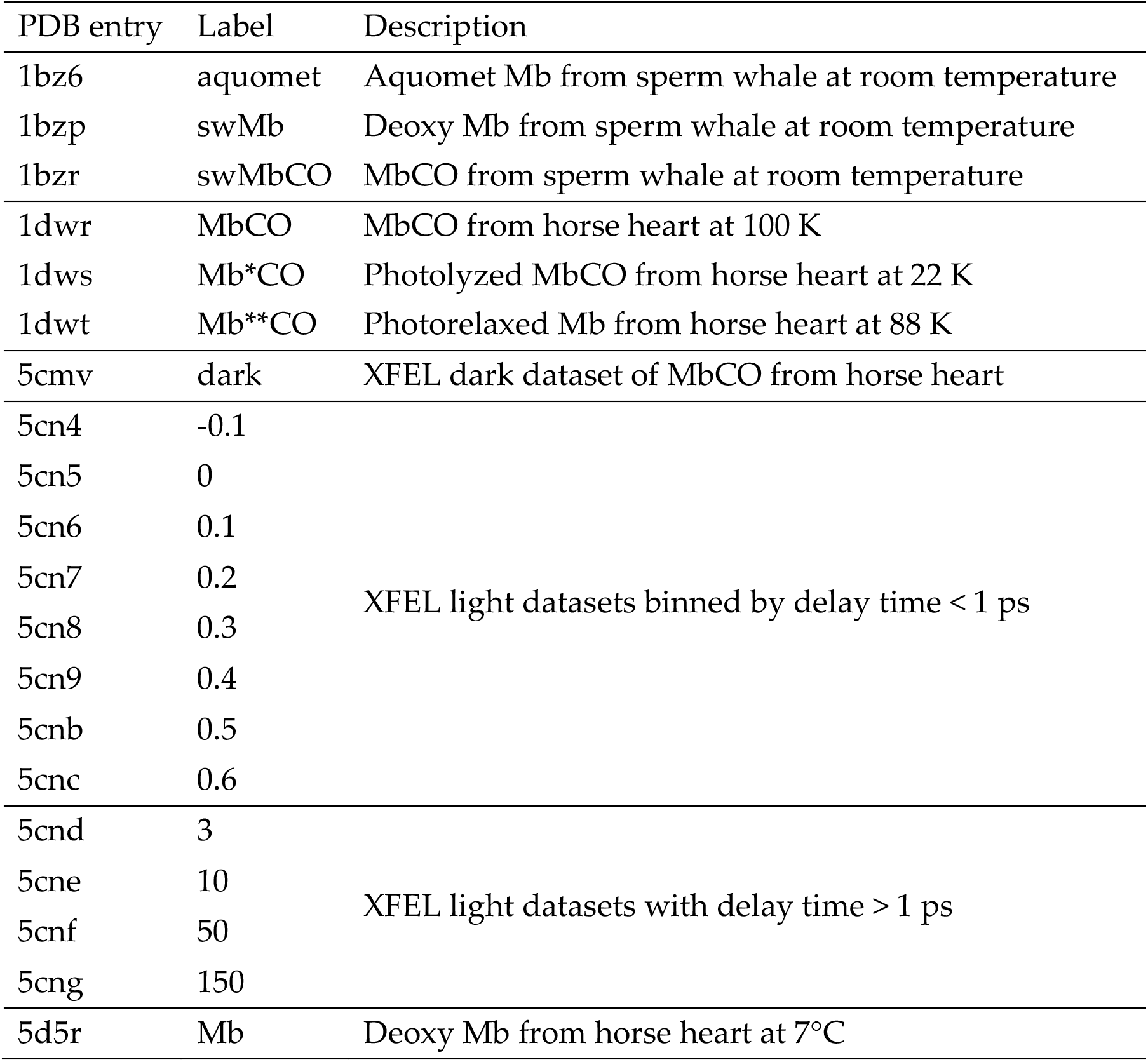
Dataset labeling

## Supplementary Figures and Legends

**Figure S1.**
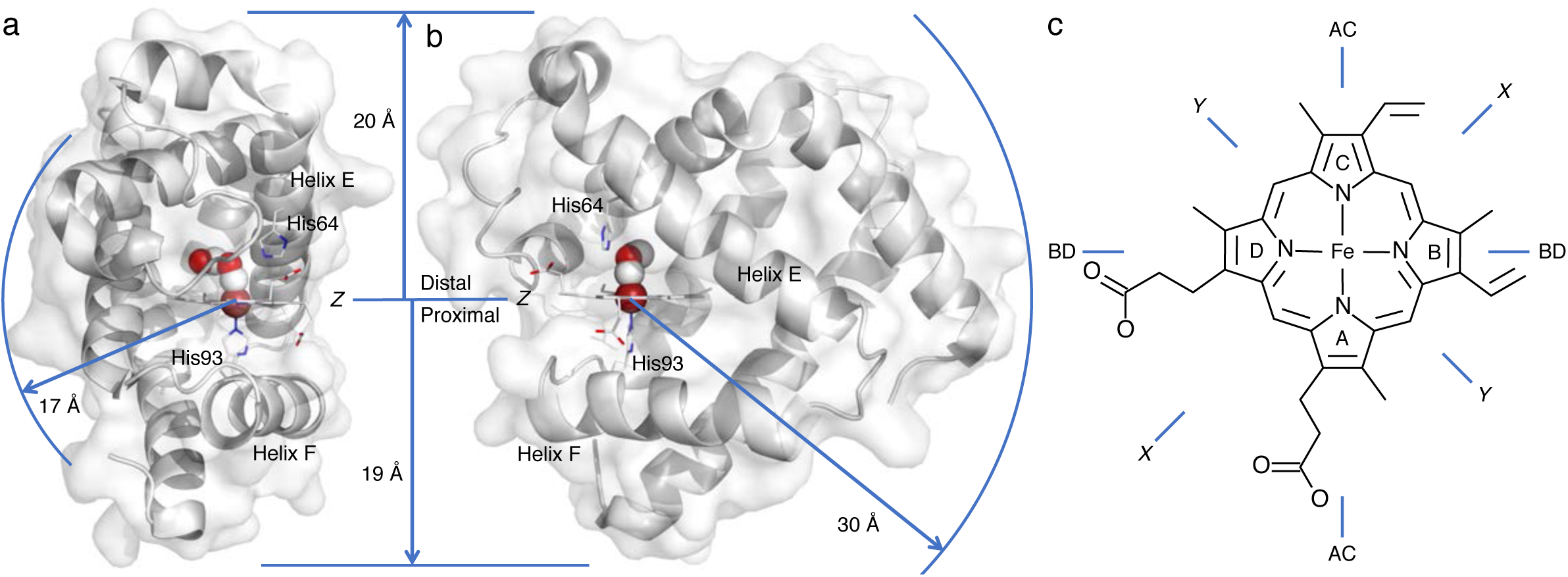
Orientations in myoglobin. Myoglobin (Mb) is a 17 kDa protein in muscle that carries molecular oxygen O_2_. The protein structure is rendered in ribbon and surface models in (a) and (b). The non-protein prosthetic group heme in stick model is anchored to helix F of the protein through His93 also in sticks. Two propionic acid side chains of the heme are facing the molecular surface. The sixth coordination site of the heme iron in sphere could bind O_2_, CO, and NO ligands also in spheres. Both a bound CO and a dissociated CO docked nearby are shown. Traditionally, the anchor side of the heme is called proximal with respect to the opposite distal side, where the ligands bind. The most remote part of the protein is 30 Å away from the heme iron. The proximal and distal directions extend 19 and 20 Å in the protein from the heme iron, respectively. The tetrapyrrole rings are traditionally named ABCD as marked in (c). In this paper, the heme plane is called *Z* cross section. Other cross sections *X*, *Y*, AC, and BD are defined in (c).

**Figure S2.**
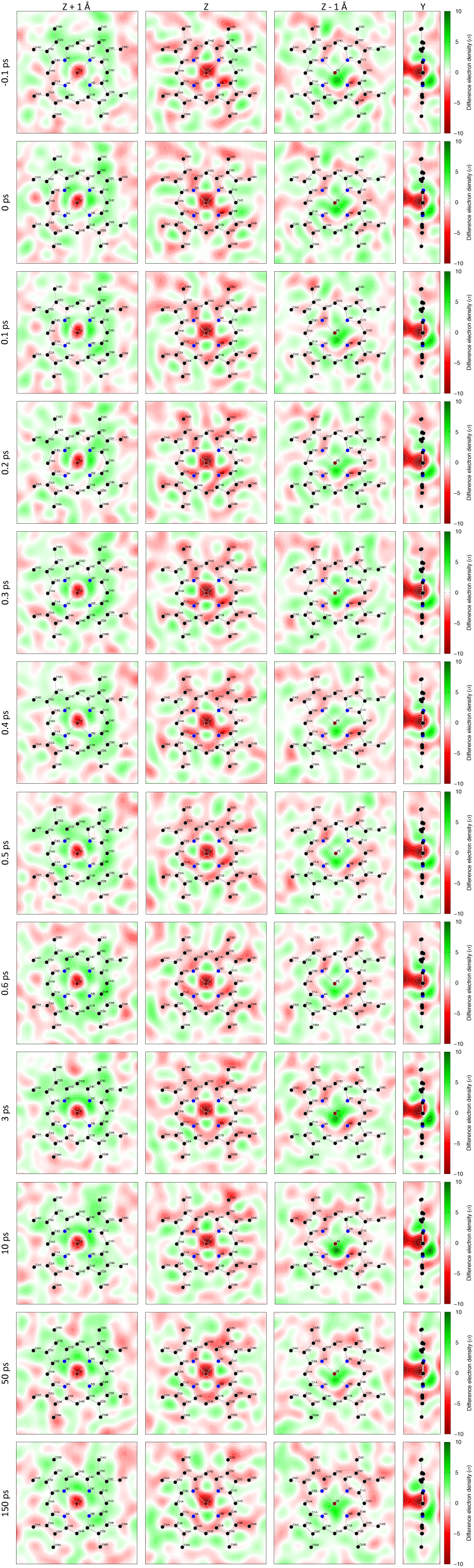
The difference Fourier maps for SVD analysis. Difference maps are calculated according to the protocol described in Methods. Only the series from −0.1 to 150 ps with the dark dataset (5cmv) as the reference is displayed here. Cross sections are marked on top. See the legends of Figs. S1 and 3 for definition of cross sections. See Table S1 for the full matrix of difference maps. A visual inspection of these difference maps at different delay times may conclude that they are largely identical. Very subtle variations could be seen by a careful comparison of these maps in a time series. A numerical deconvolution is necessary to extract the inherent components.

**Figure S3.**
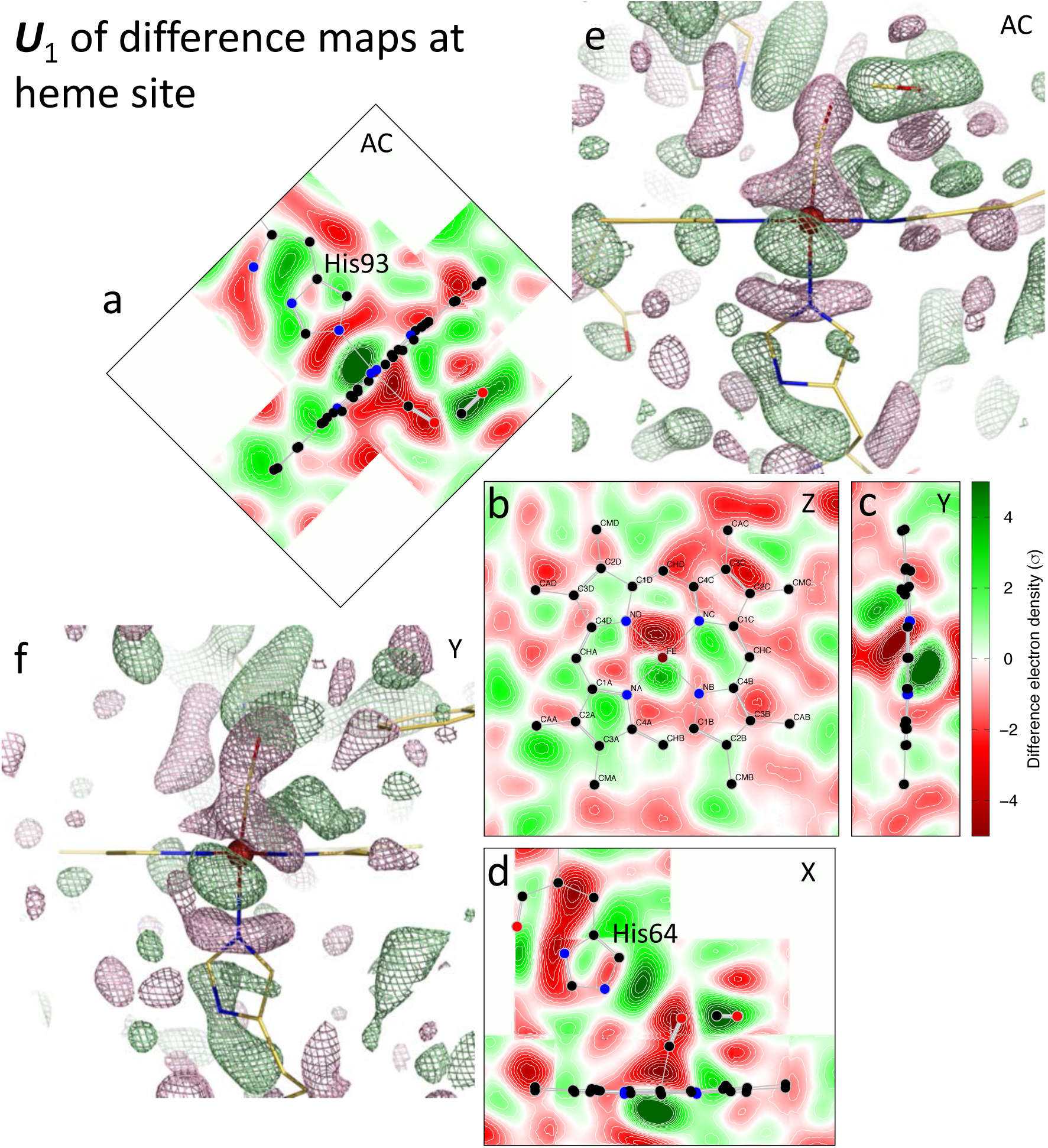
The most significant component ***U***_1_ of the difference maps. No doubt the most significant event shown in the difference maps is the out-of-plane displacement of the iron. This component shows the severely skewed iron displacement. The corresponding coefficient *c*_1_ is plotted in Fig. 2ac. This component clearly separates the delays > 1 ps from those < 1 ps. (a-d) Images of two-dimensional cross sections through the first component. (e) and (f) Three-dimensional mesh contours of the component difference map. Green and red meshes are positive and negative contours at ±7*σ*, respectively.

**Figure S4.**
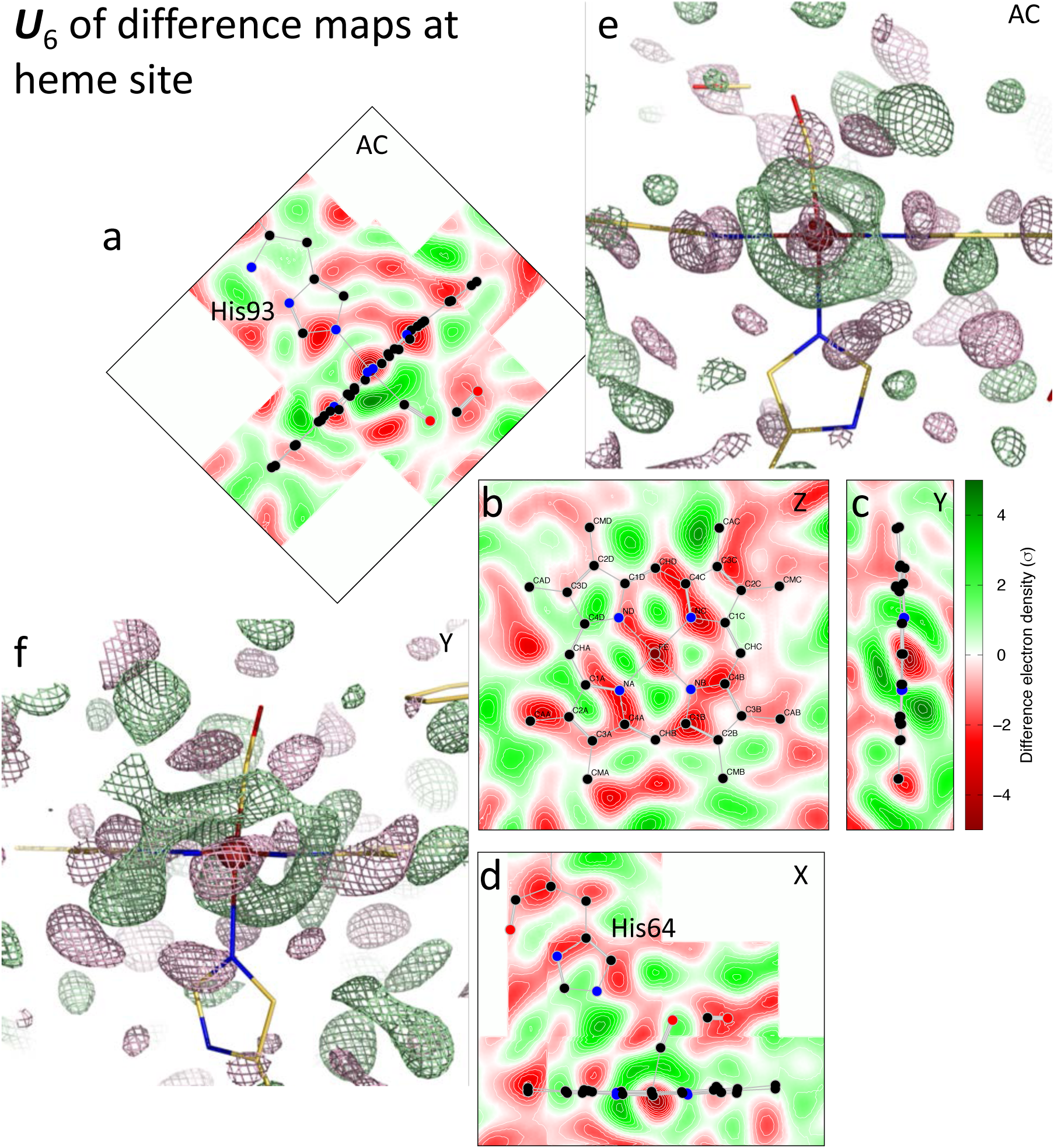
Modification to the cubic positive densities around the iron in the sixth component ***U***_6_ of the difference maps. The corresponding coefficient *c*_6_ is plotted in Fig. 2f. This component and the eighth component ***U***_8_ form a two-dimensional oscillation that ramps up in the delays > 1 ps (Fig. 2f). The effect of modulation and oscillation is shown in Movie S1. (a-d) Images of two-dimensional cross sections through the sixth component. (e) and (f) Three-dimensional mesh contours of the component difference map. Green and red meshes are positive and negative contours at ±7*σ*, respectively.

**Figure S5.**
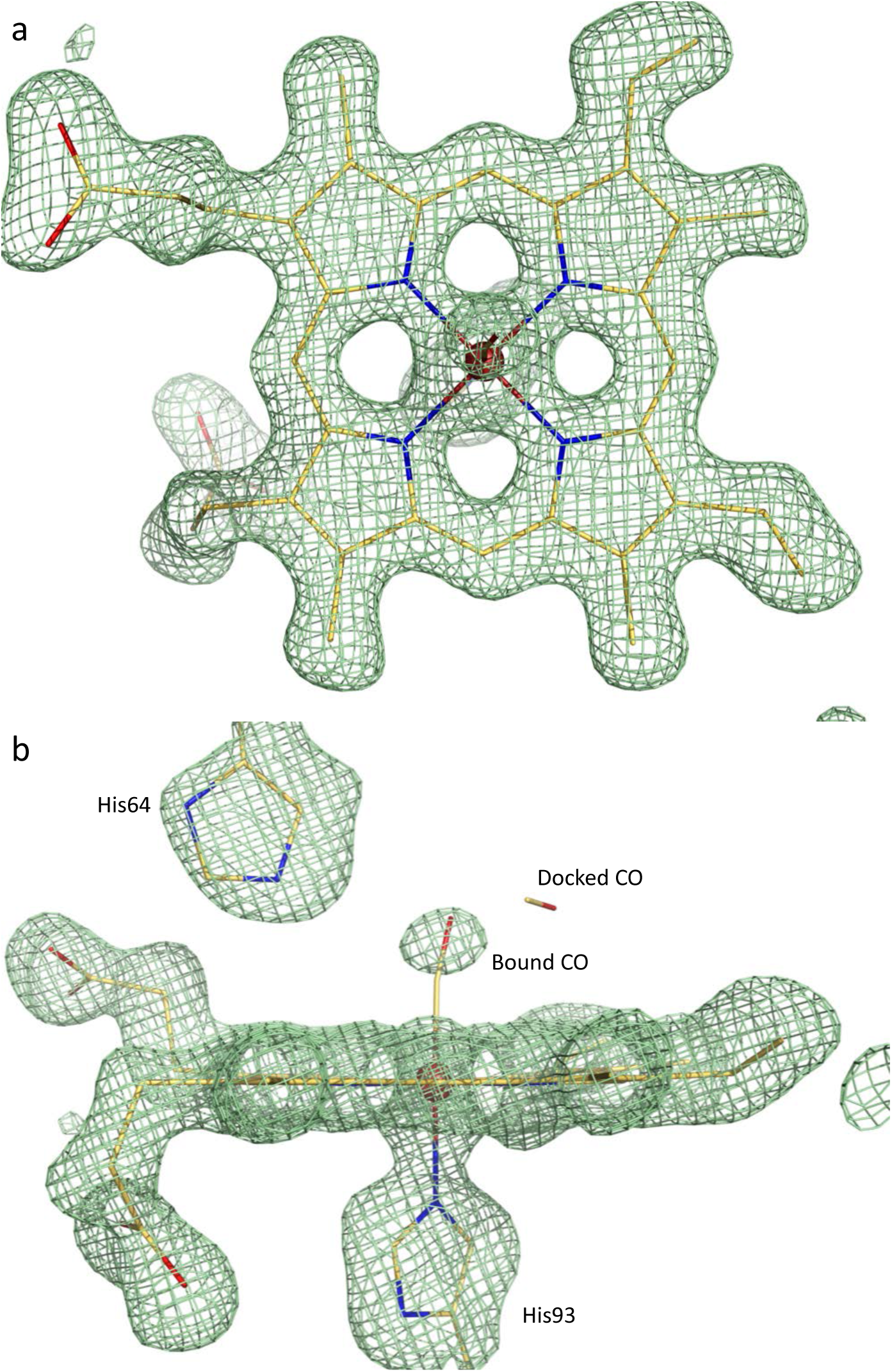
The most significant component of SAOMs. Since an omit map is the least biased experimental electron density map, the most significant component of the omit maps, by far (Fig. 4f), is an averaged experimental electron density map of 20 omit maps. Only the ligand is not clear due to its variation. All other components will be added onto or subtracted from this average.

**Movie S1.**
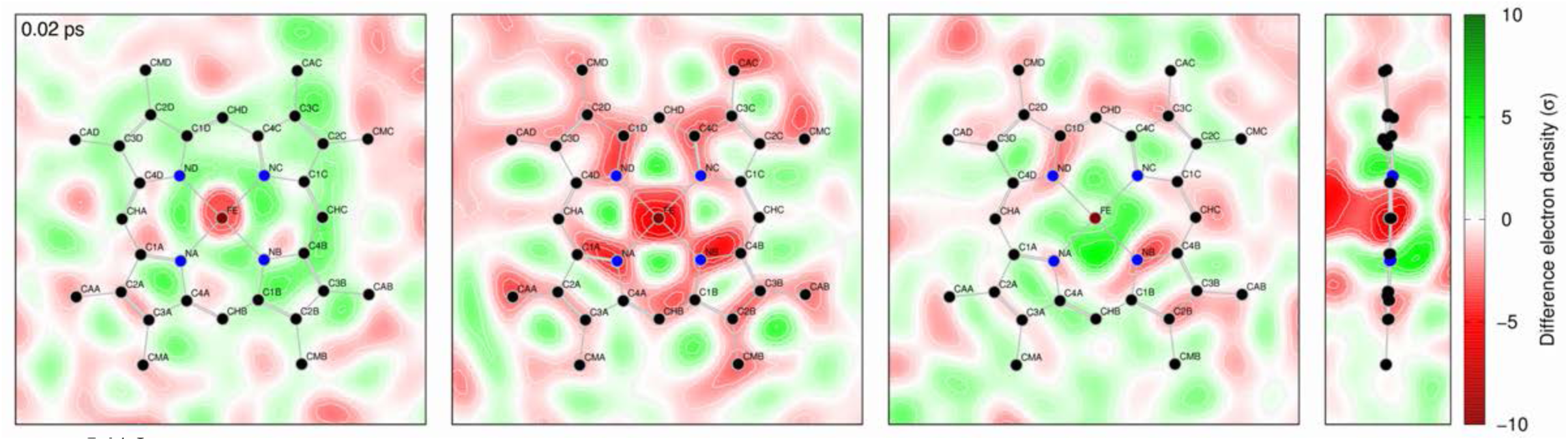
Reconstituted difference maps showing modulation and oscillation. Linear combination of components ***U***_1_, ***U***_4_, ***U***_6_, and ***U***_8_ of the difference maps span from 0.02 to 500 ps with some extrapolation on both ends. Other major components carrying systematic difference from the deoxy Mb or featuring oscillations in the solvent are not included in the linear combination to produce this mivie. The four panels are cross sections of *Z*+1.1 Å, *Z*, *Z*-1.1 Å, and *Y*, respectively. See Figs. S1 and 3 for the definition of cross sections. The cubic positive densities first lean towards ring D from 0.5 to 3 ps. Then the position returns back to the mean position after 50 ps, sways in the opposite direction towards ring B, and ramps up after 1 ps into an oscillation of a period of 220 ps (Fig. 2f). It is clear from this movie that the iron reaches its most out-of-plane displacement at a few ps and recoils back during 100-200 ps and pops out again after 200 ps.

**Movie S2.**
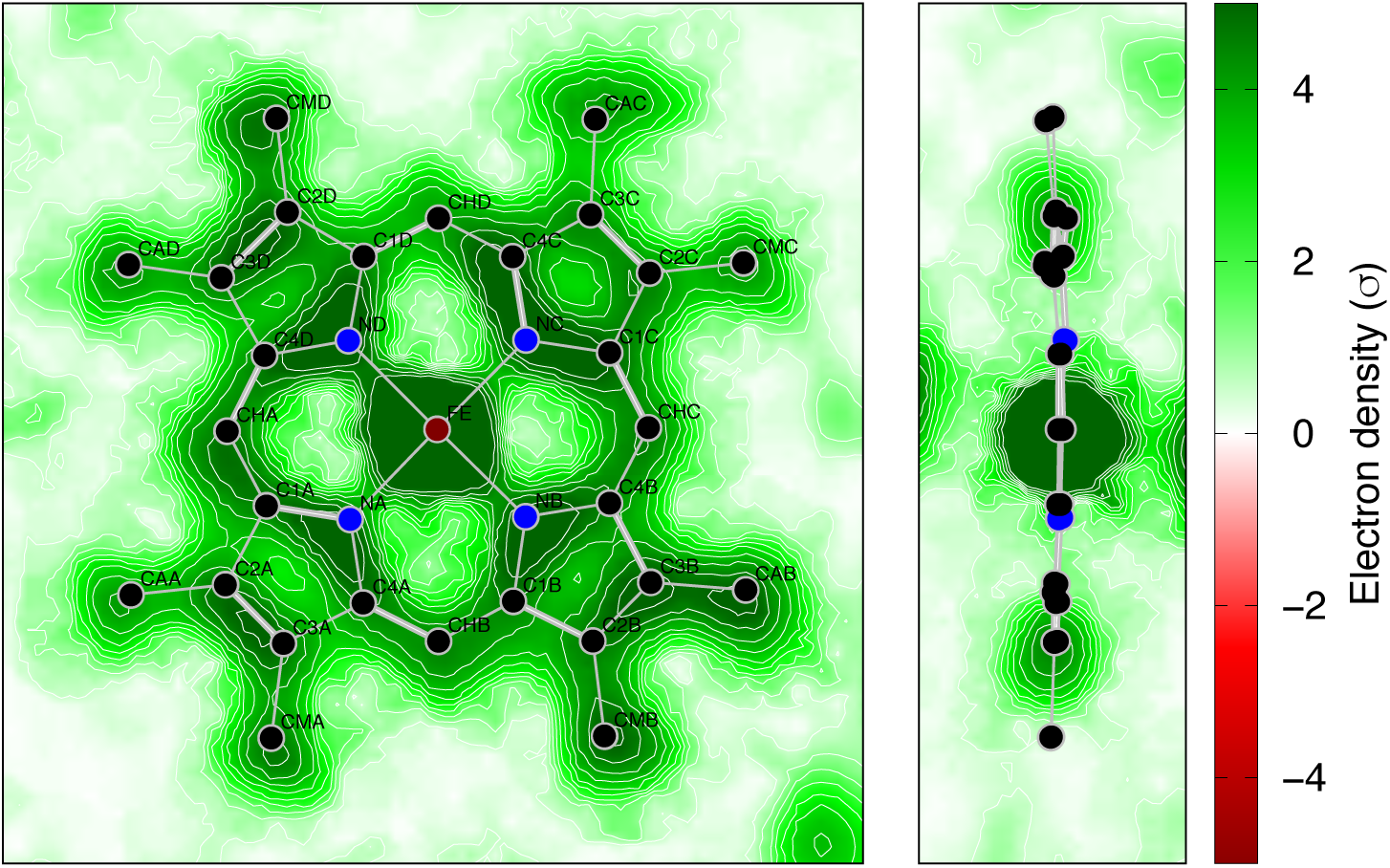
Temperature effect on electron density map of the heme. The first and third components of the SAOMs are recombined to show the temperature effect only. The components related to the chemical signals of the photolysis are not included. Therefore, this movie shows the pure thermal effect. Slight extrapolation at both ends covers from 0 to 600 K. The two panels are cross sections *Z* and *Y*, respectively. See also Fig. 6ab for electron density maps at 0 and 600 K.

**Movie S3.**
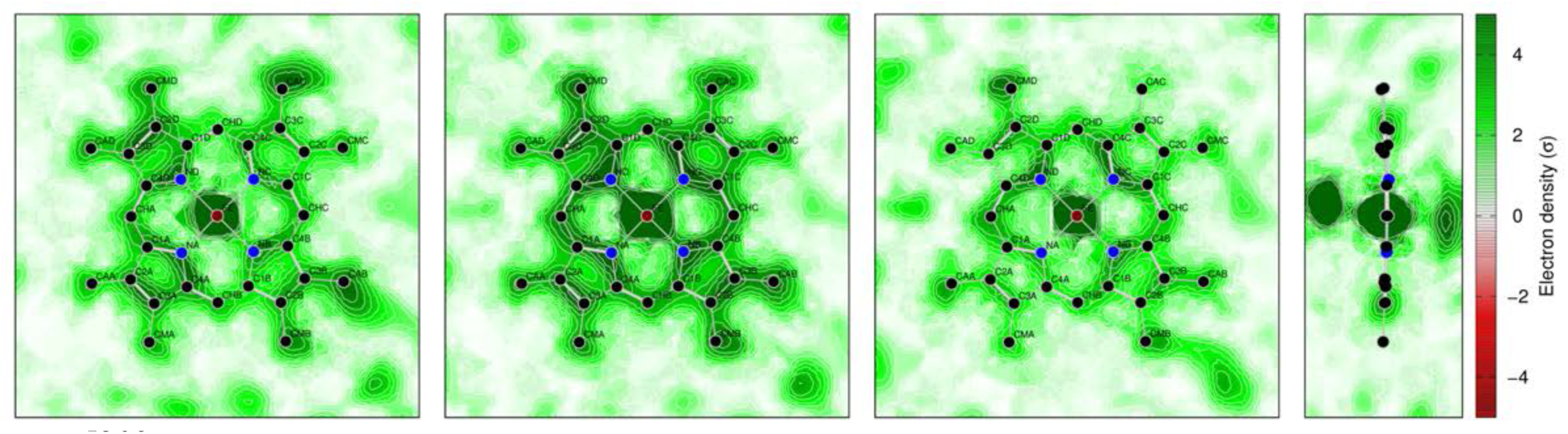
Comparison of electron density of the heme in ligated and unligated deoxy states. The first, third, and seventh components of the SAOMs are recombined to directly contrast the electron density maps of the heme in ligated and deoxy states. The coefficients for the first and third components are constant in the reconstituted maps at 0 K, that is to say, the thermal effect is excluded. The coefficient *c*_7_ for the seventh components is set to −400 for the ligated state and +300 for the deoxy state (Fig. 4e). The panels are cross sections of *Z*+1.1 Å, *Z*, *Z*-1.1 Å, and *Y*, respectively. The ligation state and the iron out-of-plane motion are shown in cross section *Y*. Cross section *Z*+1.1 Å on the distal side clearly shows greater densities of the heme in deoxy state, signaling the heme doming. Most importantly, the heme is sliding towards the AB rings in deoxy state. This is consistent with the skewed displacement of the iron and the motion of the proximal His93.

## Point-by-point responses to the reviewers’ comments

This manuscript has been extensively reviewed by two reviewers, both of whom evaluated the potential significance of the work and constructively suggested possible improvements. The reviewer 1 identified himself as Vladimir Y. Lunin of Russian Academy of Sciences. The reviewers’ comments are copied below verbatim in *italics* followed by my response. The topics are arranged by relevance to one another. The complete comments are appended at the end.

> Reviewer 1: The author challenges an ambitious task to extract tiny structural information possessing X-ray scattering data sets of a resolution only about 1.8 Å while usually this requires data of much higher resolution, atomic 1.0-1.2 Å or even ultrahigh, about 0.8 Å.
>
> Reviewer 1: In particular, this leads to the fact that individual electrons and hydrogen atoms become distinguishable in Fourier difference syntheses only when working with data sets at resolution of 1.0-1.2 Å (atomic resolution), while partial charges (density of valence electrons) become visible only at about 0.8 Å (ultra-high resolution) and only under certain restrictions on the structure refinement protocol (Afonine et al., 2004, Acta Cryst. D60, 260-274).
>
> Reviewer 2: C. Finally, some aspects of the central results could be presented with greater caution, considering that they are somewhat speculative, and that they depend on interpretation.
>
> 1) The author explains that the initial report of the observation of 3d orbitals (Zuo el al 1999) was somewhat controversial. In particular Scerri (2000) states that orbitals of single electrons are fundamentally not observable, and recommends that observations be described in terms of electron density. This would be the most cautious language. However, haven’t the philosophical arguments now run their course after 20 years, so it should now be safe to speak of the U4 component as revealing the high-spin orbitals? Unfortunately, the author’s own description shows that the electron density is only marginally interpretable as a 3d orbital. Line 368 states that the 3d orbital for an isolated Fe(II) has 0.36 Angstrom radius, and line 369 that the orbital density in an Fe4S4 cluster has a maximum radius of 1 Angstrom. Meanwhile, the data shown here place the electron density at 1.7 to 1.9 Angstrom from the iron, depending on which map is considered, so it appears that this analysis is “first of its kind” result, rather than the observation of a 3d orbital by established methods. It is somewhat difficult to understand the author’s point of view in the following text (lines 372-394). It is stated that the cubic density is not the probability distribution of a 3d electron, yet the paper states in many places that we observe a high-spin 3d orbital.

Can crystallographic data at the resolution of 1.8 Å reveal the electron density map of a single orbital? Clearly, this becomes one of the central questions to be addressed. I have consolidated previously scattered discussion into one new section in Discussion with a subtitle of “Are atomic orbitals observed?” (lines 635-710). This section should clarify my answer. A simple answer is No. However, the reasons behind this negative answer must be examined. More importantly, an accurate orbital density map of a single electron is not a goal in protein crystallography. These are my main points: First, the reason why an ultrahigh spatial resolution is required in charge-density analysis is because of the overwhelming spherical distribution of the inner electrons. Many techniques other than crystallography can capture images of atomic and molecular orbitals because these techniques are selectively sensitive to certain orbitals but not because they can achieve ultrahigh spatial resolution. If SVD is applied to charge-density analysis, the requirement for spatial resolution will reduce. Sadly, my conversation with Philip Coppens cannot continue after his pass. Second, if there is no discontinuity in any probability distribution functions of electrons, why ultrahigh frequency terms are required in Fourier synthesis? There is probably no discontinuity in the first derivatives of these distributions either, which means no kinks. Ultrahigh spatial resolution is to describe discontinuities and kinks. Therefore, ultrahigh spatial resolution appears necessary more because of the demand to high accuracy of measurements and less because of the high spatial frequencies. Third, although difference Fourier technique has been used for decades, it did not ease the demand for ultrahigh spatial resolution because conformational changes brought even more complexity into the picture, which has prevented aspherical orbital densities from showing up. What makes the difference in revealing the remanence of individual orbitals at a given median resolution is the decomposition and deconvolution (continued below).

> Reviewer 1: Creation of such tools would be a great step in development of X-ray research methods. Unfortunately, the results presented do not allow one to agree unconditionally with the author’s conclusions about the progress achieved, as well as with the interpretation of the results obtained when processing previously published experimental data. In particular, the annotated author’s conclusions on the visualization of the 3d orbitals and structural rearrangements in the spin crossover process raise significant doubts. The suggestions and findings of the author could be a basis for further discussions, but the categorical style of interpretation of the results does not correspond to the level of evidence of these conclusions and may give rise to false illusions for non-experts in X-ray data processing. All this requires a substantial revision of the manuscript and of the conception.
>
> Reviewer 2: The author is completely correct (lines 237-239) to describe the literature-reported electron densities with unique shapes as being directly influenced by electrons occupying specific orbitals. However, there is a very large difference between the current diffraction dataset that diffracts to 1.8 Angstrom resolution, and the literature report of Hirano et al. (2016), showing iron 3d and sulfur 3p density at 0.48 Angstrom resolution. It is important to specifically compare the numerical resolution limits for the Hirano and the present work at both lines 236 and 370; in order to fully communicate that the present work is not simply a routine report of orbital density. Rather, the current work is lower-resolution and more speculative in nature.

It has been stated multiple times in the revised manuscript that the captured image is NOT an accurate depiction of the 3*d* orbitals (lines 387-388 and 703-704), unlike the iron 3*d* and sulfur 3*p* orbitals at 0.48 Å. The difference in spatial resolution has been mentioned everywhere it can (lines 250 and 639). It is not a proper task for protein crystallography to provide accurate depictions of orbital densities. Instead, any direct evidence, at a good spatial resolution, to indicate a specific electronic event would be a desirable probe into the essence of biochemistry. This is one of the contributions of this work (continued below).

> Reviewer 2: The analysis leads to several astonishing claims highly relevant to myoglobin/CO binding, including that 1) the high-spin 3d orbitals can be visualized throughout the time series, 2) the XFEL pulse heats up the specimen to ∼500K, and 3) there is an oscillation with a 220 ps period.

In addition to the visual evidence to indicate the reappearance of high-spin 3*d* electrons, other conclusions such as the positional modulation, recoiling of the heme iron, the XFEL pulse induced temperature jump, and the low frequency oscillation in the solvent are as novel as the indication of the high-pin orbitals. These results are long awaited visual evidences to validate and to interpret decades of data in spectroscopy and computation. This work represents the highest achievement in serial femtosecond crystallography using XFELs. Each of the conclusions comes from a solid mathematical deduction of the XFEL datasets and corresponds to spectroscopic experiments and computational simulations in the field. However, making a reader “unconditionally” agree with my interpretations is beyond my paygrade.

> Reviewer 1: Similarly, the author ignores that, when an electron transfers from one orbital to another, the image of the orbital in a difference map should appear twice: as a negative density of the orbital that has lost an electron, and a positive density of the orbitals that have gained an electron. He does not mention either the question on a principal possibility to see such a subtle change in 1.8 Å resolution maps. The absence of the U4 component in the light-light series of difference maps is presented as evidence of the speed of this transition.

This is a very constructive suggestion. Indeed, it would be ideal to see some negative densities showing the loss of the low-spin electrons in the ligated state. However, the loss of the CO ligand and the drop of the iron cause the displacement of many more electrons than the low-spin electrons. Negative densities in the component ***U***_4_ are overwhelming. At this resolution, the loss of the low-spin electrons cannot be separated from the main events upon photolysis. This note has been added (lines 400-405).

> Reviewer 2: Investigating the light-triggered release of CO from myoglobin, the author collectively re-examines electron density maps from room temperature and cryo-trapped structures, as well as the time-dependent XFEL paper [Barends et al. (2015)] covering the −0.1 ps to 150 ps time evolution. Mutual difference maps and single-dataset omit maps are analyzed with the Schmidt et al. (2003) method of singular value decomposition (SVD), to identify features in the electron density that change with either time, sample, or temperature. This reviewer does not know if SVD has ever been applied over so many independent variables, and if so it is worth mentioning.
>
> Reviewer 2: 3) It is further stated (lines 390-393) that normal 2Fo-Fc maps cannot reveal small orbital changes, but that SVD analysis is highly sensitive. If true, this would be a critical factor, and everyone should be using SVD to analyze structure comparisons. It would be highly beneficial to lay out the evidence and controls, or provide literature references.

Applications of SVD in protein crystallography have had limited success in the past 20 years until Givens rotations are applied, mainly by me (Ren PLoS One 8, e77141 & e77363, 2013; Ren Nucleic Acids Res 6, 1, 2016). This is due to the fact that the default solution of SVD guarantees no physical meaning in the basis components. Instead, each component is a linear combination or weighted sum of several meaningful images. SVD guarantees all basis components are orthonormal. However, physically meaningful images are not necessarily orthonormal. Therefore, a satisfactory interpretation of an SVD result is based on luck, which is the “drawback” of SVD.

> Reviewer 1: An essential drawback of the suggested approach is the stage of the “manual” rotation of the found set of basic density distributions with unclear criteria of the best choice. This makes it difficult to reproduce the presented results.
>
> Reviewer 1: The author’s approach to the study of a series of experimental electron density distributions close to each other is based on a representation of each of these distributions as a weighted sum of a fixed set of density distributions pairwise uncorrelated, considered as a set of basis vectors mutually orthogonal. A visual study of the evolution of electron density under changing of external conditions is substituted by a numeric analysis of the weights with which these basis functions enter into one or another experimental density distribution. Sets of basic vectors can be selected in many ways and the basic distributions themselves may have no structural meaning. The recommendation of the author is to choose the basis in which “… an appropriate perspective can be found to elucidate the relationship between the core data and metadata clearly and concisely”. This recommendation is extremely vague, opening up the scope for the selection of the “perspective” that confirms most the hypothesis of the researcher, and makes it difficult to reproduce and validate the results obtained.
>
> Reviewer 2: 4) The left-singular vectors are not unique—this is correctly described in the supplementary material starting at line 1201. At 1249 it is stated that a Givens rotation is applied so that an appropriate perspective can be found to elucidate the relationship between the core data and metadata. However, this leaves us with a situation where the main results of the paper (e.g., the U4 difference map showing high-spin density) are not based on a deterministic procedure! To paraphrase what I understand from this description, U4 was chosen by manually finding a Givens rotation that most clearly correlates with time-dependence. However, the author does not give a protocol to find that rotation, nor a metric to judge when the rotation is optimized.

The concept of rotation was introduced much earlier by Henry & Hofrichter (in Num Comp Meth 210, 129, 1992). But their protocol specializes in the field of spectroscopy. More importantly, it fails to preserve the orthonormal and least-squares properties of SVD after rotation. This manuscript is the first time that I explicitly describe a rotation procedure with the orthonormal and least-squares properties preserved. That is to say, the rotation procedure finds an alternative solution for SVD. The rotation does not change the internal relationship among datasets except that it finds a different perspective to view the relationship. This rotation procedure may require a user decision. But this is not a drawback; this is a major advance in data analysis of dynamic crystallography. It is this rotation that finally connects the core crystallographic data to the metadata of experimental conditions. This rotation finally accomplishes the deconvolution of physical or chemical factors after decomposition. It is absolutely true that “everyone should be using SVD to analyze structure comparisons.” I also believe that this advance is not limited to crystallography. It is going to impact other fields using SVD.

For example in Fig. 2a, there is a large gap between all datasets < 1 ps and those > 1 ps, where “large” is with respect to the spacings among the datasets in each group. A rotation does not change this fact except that this gap was not originally along the first dimension *c*_1_. Therefore, the original ***U***_1_ cannot describe the structural meaning of this gap, that is, the difference between the short delays < 1 ps and the longer delays > 1 ps, because the original ***U***_1_ used to be a linear combination of several factors. The criteria of the best choice are clear, that is, to separate these two groups along a major dimension, a deterministic solution. However, did I know this large gap before SVD? Of course not. This is where a user input is necessary. Nevertheless, one cannot create this large gap by any rotation. It always exists even I don’t see it before a proper rotation is found. In fact, an interactive 3D plotting on the computer screen easily shows this large gap. What if another person decides to rotate in a different way? It is entirely possible that this large gap ends up orienting along another dimension. But the conclusion would not change. What if one decides not to apply any rotation? A linear combination of several components can still explain the inclined large gap with an identical conclusion.

For another example in Fig. 2cd, all difference maps have *c*_4_ ≍ 0 except those calculated with the dark dataset as a reference, the red trace. The criteria of the best rotation are very clear and deterministic so that ***U***_4_ depicts features only in the difference maps with the dark dataset as a reference. Again, the fact cannot be changed with any rotation that light-dark difference maps standout from the other maps except that the other maps were not originally on the plane of *c*_4_ = 0. Therefore, the original ***U***_4_ was puzzling with several physical meanings fixed in it. In this sense, it can be stated that the Ren rotation protocol completes a deconvolution of physical and chemical factors after a singular value decomposition. A more concise version of these discussions has been incorporated into Methods (lines 1365-1391).

> Reviewer 2: If I were to perform an SVD analysis on a new protein system, how would I go about applying the Ren methodology to this new system? The core issue here is reproducibility—it would be good if more details were given about how the rotations were derived, for this particular work. Would the conclusions have been robust if slightly different U-vectors were chosen?

A satisfactory interpretation in an SVD analysis, the rotation after SVD in particular, highly depends on the scientific question. A major basis component may very likely describe an uninteresting fact that has been known for many years, the so-called trivial component. A major basis may also describe a systematic error, interesting but off-topic. The key to a proper rotation is the correlation between the core data and metadata, the reason why the name meta-analysis (Ren et al. PNAS 109, 107, 2012). With a clear scientific question in mind, a rotation is deterministic. However, before the data are evaluated by SVD, it is indeed difficult to predict what SVD may turn out. Since two different physical parameters, say temperature and pH, may not cause orthogonal changes to a structure, a single rotation is not sufficient to satisfy both dimensions. That is to say, multiple rotations, thus multiple perspectives, are needed to answer all the scientific questions. If two parameters are largely orthogonal to each other, the solutions are deterministic and stable. However, if two parameters are highly colinear, that is, they cause very similar structural responses, the solutions could be less stable. Discussion and Methods have been expanded. However, this is not a paper on applications of SVD.

> Reviewer 1: A key problem in identifying small changes in the electron density of an object in a series of experiments under variable conditions is that one never has the true distribution of electron density. What a researcher has instead is a Fourier synthesis of electron density, always of a limited resolution. Saying nothing about experimental errors, these syntheses contain larger or smaller distortion, caused by lack of the necessary data. The magnitude of such distortions, called Fourier series truncation errors, can be significant and these distortions can mask the subtle changes in the electron density.
>
> Reviewer 1: Even 0.6 Å-resolution syntheses can contain Fourier series truncation errors exceeding the signal level and mimicking structural details. The results presented by the author have been obtained at resolution of 1.8 Å, which is intermediate between “medium” and “high” resolutions according to the existing crystallographic classification. At such conditions, sufficiently large distortions of Fourier syntheses are expected. In this regard, the questions arise: “What are the changes studied in the manuscript? Are they those in the electronic structure of the sample or just a game of the Fourier series truncation errors?” The author does not pay any attention to these circumstances. Moreover, the author regularly speaks of “ultrahigh resolution”, which can confuse the reader.

The topic of Fourier truncation error is not new. Its impact to electron density maps has been well understood. Errors from this source are not an unpredictable ghost. Therefore, we cannot attribute everything that we don’t understand to this error. First, a Fourier truncation error may most likely occur to several static datasets included in this analysis, in which the spatial resolutions are truncated to 1.75 Å to match that of XFEL datasets. However, such errors are not clearly present in these datasets, thanks to the difference Fourier technique and the simulated annealing protocol. Second, when the average intensity of an experimental dataset naturally decays as function of the spatial resolution, Fourier truncation errors are damped to a minimum. A low resolution dataset not necessarily contains more severe Fourier truncation errors; it simply lacks more details. Third, difference Fourier technique again self-cancels the remaining errors because both datasets would carry similar Fourier truncation errors. Fourth, the applied weighting scheme severely reduces any large differences in Fourier synthesis (Methods), which also prevents Fourier truncation errors. Fifth, even when Fourier truncation errors occur, such errors may appear as minor components after SVD (Figs. 2e and 4f), therefore isolated from the signals under discussion. Sixth, the SVD rotations validate that the signals under discussion, such as the positive cubic network, is not due to Fourier truncation errors because of its correlation with the metadata. An artifact due to Fourier truncation errors, if so conclude, needs to be explained why it is correlated with a physical or chemical parameter. As a conclusion, Fourier truncation errors are well guarded from. New findings cannot be dismissed under the fear of such errors. Discussion on Fourier truncation errors has been added (lines 679-695).

> Reviewer 1: The start data is a set of several series of difference maps. The first series is based on the differences between results of the “dark” (without the use of exciting light pulses) experiment and ones in which the crystals were irradiated both light X-ray pulses (“light” sets), with different time delay F_i^light-k_(i,0)·F_^dark, i=1,2,3,… The remaining difference maps were calculated using only “light” data sets F_i^light-k_(i,j)·F_j^light, i=j,j+1,… Here k_(i,j) are scaling factors, individual for each map. I start with a naive crystallographic explanation of the results. The first series of maps is shown in Fig. S2. These maps do not reproduce the typical pattern of difference maps, but rather reproduce a flipped (the one with the sign changed) image of the electron density distribution in the vicinity of the heme (the initial distribution is shown in Fig. S5). This is apparently caused by a imbalance in the difference syntheses of the first series, i.e. by too large values of the coefficients k_(i,0), that results in the -F^dark component dominating the maps.

It remains very puzzling to me after many readings of the above comments that Lunin, the first reviewer, calls the difference maps displayed in Fig. S2 “not typical” and “flipped sign”. There are two ways to interpret Lunin’s comments: The positive and negative densities in the difference maps are mistakenly flipped or the difference maps look like a negation of an electron density map in Fig. S5. Neither makes sense to me. The convention in producing these difference maps is light-dark so that positive densities show the product of light and negative densities show the loss of the dark state. The first column of Fig. S2 Z+1Å shows a cross section parallel to the heme plane and 1 Å towards to distal side. The second column Z shows the heme plane. It is obvious that Z+1Å cross section has more positive densities and Z cross section has more negative densities. This is a typical difference map of light-dark at the heme site and cannot have a flipped sign because the heme buckles towards the distal side as a result of the light illumination. Also, the bound CO ligand is in an intense negative peak indicating the loss of the ligand and an intense positive peak skewed on the proximal side of the iron (right column in Fig. S2). Again, this is typical and without sign flipping. These maps are identical to those originally produced by Barends & Schlichting (Science 350, 445, 2015) except in an arguably better quality. The scaling of the light and dark datasets is perfectly balanced as judged by equal amounts of positive and negative densities. Alternatively, a sign flipped image from Fig. S5 in Fig. S2? Of course, this is light-dark, the part of -dark should be a sign flipped image (see below). It seems that there exist some fundamental misunderstandings I have not figured out.

> Reviewer 1: After decomposition, a balanced differential density distribution appears as the basic U1 distribution (Fig. S3). This map is close to the difference map of 1.8 Å resolution. The excessive presence of the term -F^dark in the initial difference syntheses F^light-[k·F] ^dark is compensated by U4 component (Fig. 3c), which resembles again the flipped image of density in the vicinity of the heme (Fig. S5). As noted by the author, in light-to-light difference syntheses F_i^light-k·F_j^light the U4 component is absent. This can be explained by a better balance of syntheses coefficients when working with data sets obtained under the same experimental conditions (in the presence of both “light” and “dark” X-ray pulses) and, as a result, the absence of a “ghost” corresponding to the initial heme density distribution.

Essentially, Lunin questions a possibility that the light-dark series is not scaled as well as the other light-light series, because the light and dark datasets were obtained under slightly different conditions, such as no laser pulse for the dark dataset. As a result, a “ghost” component exists in the light-dark series but not in the light-light series that has been decomposed by SVD as ***U***_4_. The principle in meta-analysis is the correlation between core data and metadata. If the component ***U***_4_ is an artifact due to scaling errors, it must be explained why these errors remain constant for short delays < 1 ps and decay as function of time > 1 ps. The most significant systematic error should occur to the deoxy dataset collected at a synchrotron instead of an XFEL (marked as Mb in Fig. 2). However, this dataset fits smoothly into the trace. Again, an unusual result cannot be simply dismissed by an unexplained “error”, here a scaling error.

> Reviewer 1: The author suggests a totally different explanation declaring U4 component as visualization of 3d orbital. An attention is not paid to the similarity of this distribution with the density distributions in the vicinity of the heme, shown in Fig. S2 and S5, as well as to the discrepancy between the size of the orbitals and the size of the density fragments in the U4 component noted by the author.

Again, this comment is very puzzling to me. There is a negative image of Fig. S5 in Fig. S2. This is correct, since Fig. S2 shows light-dark difference maps and Fig. S5 shows an average map of all datasets, which is very close to the dark map. The negative part of Fig. S2 is supposed to be similar to Fig. S5. As the size of the density features, there is not enough spatial resolution to depict an accurate map of the 3*d* orbitals (see above). However, this is not to say that the net gain of densities further away from the iron is completely meaningless. The shape of the net gain coincides with the calculated orbital densities, which strongly suggests the influence of the high-spin 3*d* orbitals.

> Reviewer 2: 3) It is difficult to understand why the time evolution in Fig. 2f is described as an “oscillation”. Not even one full cycle is observed, so it should not be assumed to be a circular progression. If there were more time points available they could prove or disprove the model.

Here an oscillation is only a model that requires the least number of parameters to describe the variation. This can be updated when more observations become available (line 460).

> Reviewer 2: A. The author does a good job at describing how the analysis was done, and providing literature references to place the work in context. However, I am concerned that the data presentation and logical arguments are quite lengthy and detailed, and are challenging to keep straight. To assist the wider audience, a few things could readily be done to enhance the text’s accessibility:

This reviewer offered a number of excellent suggestions to make the manuscript better.

> Reviewer 2: 2) The claim that the XFEL beam pumps the sample to 500 K relies on evidence that is quite indirect—Fig 6i assumes that the synchrotron datasets at various thermometer temperatures can be plotted on a straight line with respect to the c3 coefficient, and that the XFEL data can then be placed directly on this line. It would be more cautious if the text admitted that there is no other physical evidence for this claim. In other words, this is an interesting but unique claim for which other evidence should be gathered.

This is a good suggestion. A note has been added to state that the temperature jump is solely based on a linear extrapolation (lines 525-527).

> Reviewer 2: B. The subject matter and techniques are extremely specialized, and I believe that more must be done to help the audience place the methods and claims into a larger context. Four specific concerns are:
>
> 1) Even for an expert protein crystallographer, the spin states of CO-myoglobin may not be common knowledge. It is important to give enough basic chemistry background in the Introduction so the reader does not erroneously conclude that the data actually distinguish between high spin and low spin electrons. Rather, we know fundamentally that in MbFe(II)CO, Fe(II) has 6 electrons, all are paired so the total spin is S=0. This is because the 5 d-orbitals are split by the ligand field into 3 lower t2g + 2 upper eg; so the lower orbitals are all full. On photolysis of CO, the ligand field and orbital splitting decrease, so there are four unpaired electrons with total spin S=2. Including this textbook material would help to capture a larger audience.

This is an excellent point. I have incorporated a little more into the brief background of spin crossover in Introduction (lines 108-116).

> Reviewer 2: 1) There are exactly 11 novel claims made in Fig. 1, represented by blue lines in the time chart with corresponding text description. It would be good for each of these blue lines to indicate which Figures contain the supporting evidence. For example, the “heme heating up to 500-600K” is clearly a consequence of Fig. 6i. Cross-referencing in this manner would help the reader keep track of the logical arguments.
>
> 2) A similar comment applies to the claims in the Introduction. Clearly line 172 “high spin iron” follows from Fig. 3; however, this reviewer cannot locate the evidence for “high frequency vibrational mode” and “low frequency oscillations” in lines 178-179.

I have implemented these cross referencing.

> Reviewer 2: 3) Part of the problem is that the reader does not necessarily have the information about what is “high frequency” and “low frequency” in a particular context; after all, the paper discusses 18-orders of magnitude of time in Fig. 1. It seems safer to avoid the use of “high” and “low” and possibly give specific numerical ranges or functional descriptions.

I have added the corresponding wavenumbers where high or low frequency is mentioned.

> Reviewer 2: 4) Also the line 213-214 claim that an oscillation occurs more in the solvent channels than in the protein or heme is similarly difficult to find in the visual evidence. Is it within Figure 6 somewhere? One possible strategy would be to take specific claims out of the Introduction since they are repeated anyway in the sequence of Results.

Fig. 7 is about the oscillation largely in the solvent. Cross referencing has been added.

> Reviewer 2: Various typographical and language corrections were found:
>
> Line 56 “of hundreds of K”
> Line 80 “first few”
> Line 151 “preemptively avoiding”
> Line 192 “not sufficient time points”
> Line 204 please specify what kind of shift
> Line 577 “features can be found”
> Line 608 “structural”
> Line 630 “conceals”
> Line 929 “first few”

All fixed.

Reviewers’ comments:

Reviewer #1 (Remarks to the Author):

The author challenges an ambitious task to extract tiny structural information possessing X-ray scattering data sets of a resolution only about 1.8 Å while usually this requires data of much higher resolution, atomic 1.0-1.2 Å or even ultrahigh, about 0.8 Å. Creation of such tools would be a great step in development of X-ray research methods. Unfortunately, the results presented do not allow one to agree unconditionally with the author’s conclusions about the progress achieved, as well as with the interpretation of the results obtained when processing previously published experimental data. In particular, the annotated author’s conclusions on the visualization of the 3d orbitals and structural rearrangements in the spin crossover process raise significant doubts. An essential drawback of the suggested approach is the stage of the “manual” rotation of the found set of basic density distributions with unclear criteria of the best choice. This makes it difficult to reproduce the presented results. The suggestions and findings of the author could be a basis for further discussions, but the categorical style of interpretation of the results does not correspond to the level of evidence of these conclusions and may give rise to false illusions for non-experts in X-ray data processing. All this requires a substantial revision of the manuscript and of the conception.

A key problem in identifying small changes in the electron density of an object in a series of experiments under variable conditions is that one never has the true distribution of electron density. What a researcher has instead is a Fourier synthesis of electron density, always of a limited resolution. Saying nothing about experimental errors, these syntheses contain larger or smaller distortion, caused by lack of the necessary data. The magnitude of such distortions, called Fourier series truncation errors, can be significant and these distortions can mask the subtle changes in the electron density. In particular, this leads to the fact that individual electrons and hydrogen atoms become distinguishable in Fourier difference syntheses only when working with data sets at resolution of 1.0-1.2 Å (atomic resolution), while partial charges (density of valence electrons) become visible only at about 0.8 Å (ultra-high resolution) and only under certain restrictions on the structure refinement protocol (Afonine et al., 2004, Acta Cryst. D60, 260-274). Even 0.6 Å-resolution syntheses can contain Fourier series truncation errors exceeding the signal level and mimicking structural details. The results presented by the author have been obtained at resolution of 1.8 Å, which is intermediate between “medium” and “high” resolutions according to the existing crystallographic classification. At such conditions, sufficiently large distortions of Fourier syntheses are expected. In this regard, the questions arise: “What are the changes studied in the manuscript? Are they those in the electronic structure of the sample or just a game of the Fourier series truncation errors?” The author does not pay any attention to these circumstances. Moreover, the author regularly speaks of “ultrahigh resolution”, which can confuse the reader.

The author’s approach to the study of a series of experimental electron density distributions close to each other is based on a representation of each of these distributions as a weighted sum of a fixed set of density distributions pairwise uncorrelated, considered as a set of basis vectors mutually orthogonal. A visual study of the evolution of electron density under changing of external conditions is substituted by a numeric analysis of the weights with which these basis functions enter into one or another experimental density distribution. Sets of basic vectors can be selected in many ways and the basic distributions themselves may have no structural meaning. The recommendation of the author is to choose the basis in which “… an appropriate perspective can be found to elucidate the relationship between the core data and metadata clearly and concisely”. This recommendation is extremely vague, opening up the scope for the selection of the “perspective” that confirms most the hypothesis of the researcher, and makes it difficult to reproduce and validate the results obtained.

The start data is a set of several series of difference maps. The first series is based on the differences between results of the “dark” (without the use of exciting light pulses) experiment and ones in which the crystals were irradiated both light X-ray pulses (“light” sets), with different time delay F_i^light-k_(i,0)·F_^dark, i=1,2,3,… The remaining difference maps were calculated using only “light” data sets F_i^light-k_(i,j)·F_j^light,i=j,j+1,… Here k_(i,j) are scaling factors, individual for each map.

I start with a naive crystallographic explanation of the results. The first series of maps is shown in Fig. S2. These maps do not reproduce the typical pattern of difference maps, but rather reproduce a flipped (the one with the sign changed) image of the electron density distribution in the vicinity of the heme (the initial distribution is shown in Fig. S5). This is apparently caused by a imbalance in the difference syntheses of the first series, i.e. by too large values of the coefficients k_(i,0), that results in the -F^dark component dominating the maps. After decomposition, a balanced differential density distribution appears as the basic U1 distribution (Fig. S3). This map is close to the difference map of 1.8 Å resolution. The excessive presence of the term -F^dark in the initial difference syntheses F^light-[k·F]^dark is compensated by U4 component (Fig. 3c), which resembles again the flipped image of density in the vicinity of the heme (Fig. S5). As noted by the author, in light-to-light difference syntheses F_i^light-k·F_j^light the U4 component is absent. This can be explained by a better balance of syntheses coefficients when working with data sets obtained under the same experimental conditions (in the presence of both “light” and “dark” X-ray pulses) and, as a result, the absence of a “ghost” corresponding to the initial heme density distribution.

The author suggests a totally different explanation declaring U4 component as visualization of 3d orbital. An attention is not paid to the similarity of this distribution with the density distributions in the vicinity of the heme, shown in Fig. S2 and S5, as well as to the discrepancy between the size of the orbitals and the size of the density fragments in the U4 component noted by the author. Similarly, the author ignores that, when an electron transfers from one orbital to another, the image of the orbital in a difference map should appear twice: as a negative density of the orbital that has lost an electron, and a positive density of the orbitals that have gained an electron. He does not mention either the question on a principal possibility to see such a subtle change in 1.8 Å resolution maps. The absence of the U4 component in the light-light series of difference maps is presented as evidence of the speed of this transition. By this particular example, I would like to illustrate that the results obtained allow different interpretations that could be a subject of discussion, and that the author’s interpretation is rather vague and should not be presented as a proof.

Vladimir Y. Lunin

Reviewer #2 (Remarks to the Author):

Investigating the light-triggered release of CO from myoglobin, the author collectively re-examines electron density maps from room temperature and cryo-trapped structures, as well as the time-dependent XFEL paper [Barends et al. (2015)] covering the −0.1 ps to 150 ps time evolution. Mutual difference maps and single-dataset omit maps are analyzed with the Schmidt et al. (2003) method of singular value decomposition (SVD), to identify features in the electron density that change with either time, sample, or temperature. This reviewer does not know if SVD has ever been applied over so many independent variables, and if so it is worth mentioning.

The analysis leads to several astonishing claims highly relevant to myoglobin/CO binding, including that 1) the high-spin 3d orbitals can be visualized throughout the time series, 2) the XFEL pulse heats up the specimen to ∼500K, and 3) there is an oscillation with a 220 ps period.

A. The author does a good job at describing how the analysis was done, and providing literature references to place the work in context. However, I am concerned that the data presentation and logical arguments are quite lengthy and detailed, and are challenging to keep straight. To assist the wider audience, a few things could readily be done to enhance the text’s accessibility:

1) There are exactly 11 novel claims made in Fig. 1, represented by blue lines in the time chart with corresponding text description. It would be good for each of these blue lines to indicate which Figures contain the supporting evidence. For example, the “heme heating up to 500-600K” is clearly a consequence of Fig. 6i. Cross-referencing in this manner would help the reader keep track of the logical arguments.

2) A similar comment applies to the claims in the Introduction. Clearly line 172 “high spin iron” follows from Fig. 3; however, this reviewer cannot locate the evidence for “high frequency vibrational mode” and “low frequency oscillations” in lines 178-179.

3) Part of the problem is that the reader does not necessarily have the information about what is “high frequency” and “low frequency” in a particular context; after all, the paper discusses 18-orders of magnitude of time in Fig. 1. It seems safer to avoid the use of “high” and “low” and possibly give specific numerical ranges or functional descriptions.

4) Also the line 213-214 claim that an oscillation occurs more in the solvent channels than in the protein or heme is similarly difficult to find in the visual evidence. Is it within Figure 6 somewhere? One possible strategy would be to take specific claims out of the Introduction since they are repeated anyway in the sequence of Results.

B. The subject matter and techniques are extremely specialized, and I believe that more must be done to help the audience place the methods and claims into a larger context. Four specific concerns are:

1) Even for an expert protein crystallographer, the spin states of CO-myoglobin may not be common knowledge. It is important to give enough basic chemistry background in the Introduction so the reader does not erroneously conclude that the data actually distinguish between high spin and low spin electrons. Rather, we know fundamentally that in MbFe(II)CO, Fe(II) has 6 electrons, all are paired so the total spin is S=0. This is because the 5 d-orbitals are split by the ligand field into 3 lower t2g + 2 upper eg; so the lower orbitals are all full. On photolysis of CO, the ligand field and orbital splitting decrease, so there are four unpaired electrons with total spin S=2. Including this textbook material would help to capture a larger audience.

2) The author is completely correct (lines 237-239) to describe the literature-reported electron densities with unique shapes as being directly influenced by electrons occupying specific orbitals. However, there is a very large difference between the current diffraction dataset that diffracts to 1.8 Angstrom resolution, and the literature report of Hirano et al. (2016), showing iron 3d and sulfur 3p density at 0.48 Angstrom resolution. It is important to specifically compare the numerical resolution limits for the Hirano and the present work at both lines 236 and 370; in order to fully communicate that the present work is not simply a routine report of orbital density. Rather, the current work is lower-resolution and more speculative in nature.

3) It is further stated (lines 390-393) that normal 2Fo-Fc maps cannot reveal small orbital changes, but that SVD analysis is highly sensitive. If true, this would be a critical factor, and everyone should be using SVD to analyze structure comparisons. It would be highly beneficial to lay out the evidence and controls, or provide literature references.

4) The left-singular vectors are not unique—this is correctly described in the supplementary material starting at line 1201. At 1249 it is stated that a Givens rotation is applied so that an appropriate perspective can be found to elucidate the relationship between the core data and metadata. However, this leaves us with a situation where the main results of the paper (e.g., the U4 difference map showing high-spin density) are not based on a deterministic procedure! To paraphrase what I understand from this description, U4 was chosen by manually finding a Givens rotation that most clearly correlates with time-dependence. However, the author does not give a protocol to find that rotation, nor a metric to judge when the rotation is optimized. If I were to perform an SVD analysis on a new protein system, how would I go about applying the Ren methodology to this new system? The core issue here is reproducibility—it would be good if more details were given about how the rotations were derived, for this particular work. Would the conclusions have been robust if slightly different U-vectors were chosen?

C. Finally, some aspects of the central results could be presented with greater caution, considering that they are somewhat speculative, and that they depend on interpretation.

1) The author explains that the initial report of the observation of 3d orbitals (Zuo el al 1999) was somewhat controversial. In particular Scerri (2000) states that orbitals of single electrons are fundamentally not observable, and recommends that observations be described in terms of electron density. This would be the most cautious language. However, haven’t the philosophical arguments now run their course after 20 years, so it should now be safe to speak of the U4 component as revealing the high-spin orbitals? Unfortunately, the author’s own description shows that the electron density is only marginally interpretable as a 3d orbital. Line 368 states that the 3d orbital for an isolated Fe(II) has 0.36 Angstrom radius, and line 369 that the orbital density in an Fe4S4 cluster has a maximum radius of 1 Angstrom. Meanwhile, the data shown here place the electron density at 1.7 to 1.9 Angstrom from the iron, depending on which map is considered, so it appears that this analysis is “first of its kind” result, rather than the observation of a 3d orbital by established methods. It is somewhat difficult to understand the author’s point of view in the following text (lines 372-394). It is stated that the cubic density is not the probability distribution of a 3d electron, yet the paper states in many places that we observe a high-spin 3d orbital.

2) The claim that the XFEL beam pumps the sample to 500 K relies on evidence that is quite indirect—Fig 6i assumes that the synchrotron datasets at various thermometer temperatures can be plotted on a straight line with respect to the c3 coefficient, and that the XFEL data can then be placed directly on this line. It would be more cautious if the text admitted that there is no other physical evidence for this claim. In other words, this is an interesting but unique claim for which other evidence should be gathered.

3) It is difficult to understand why the time evolution in Fig. 2f is described as an “oscillation”. Not even one full cycle is observed, so it should not be assumed to be a circular progression. If there were more time points available they could prove or disprove the model.

Various typographical and language corrections were found:

Line 56 “of hundreds of K”
Line 80 “first few”
Line 151 “preemptively avoiding”
Line 192 “not sufficient time points”
Line 204 please specify what kind of shift
Line 577 “features can be found”
Line 608 “structural”
Line 630 “conceals”
Line 929 “first few”

